# Seasonal Variation Mediates the Importance of Species Attributes in Plant-Pollinator Interactions

**DOI:** 10.64898/2026.07.21.739935

**Authors:** Jose B. Lanuza, Alfonso Allen-Perkins, Will Glenny, Anna Traveset, Henriette F. Morgenroth, Raymond Umazekabiri, Marc Hoffmann, Panagiotis Theodorou, Robert J. Paxton, Isabell Hensen, Robert Rauschkolb, Christine Römermann, Oliver Schweiger, Tiffany Knight

## Abstract

Predicting species interactions remains a major challenge, as multiple species attributes operate simultaneously and their relative importance may vary seasonally and with temporal resolution. Here, we assess how the relative contributions of abundance, trait matching, and phenology to plant-pollinator interactions vary through a flowering season and across temporal resolutions using interaction data from three European botanical gardens. We show that predictive models explain a substantial proportion of variation in visitation patterns, with floral and pollinator abundances consistently explaining most variation across the flowering season. Trait matching between pollinator body size and floral size also contributes to visitation patterns, playing a secondary but persistent role in shaping interactions, while phenology plays a relatively minor role at broader temporal resolutions but becomes more important at finer temporal scales. Additionally, null models accounting for spatio-temporal variation in floral and pollinator abundance reveal consistent patterns of pollinator preference and avoidance, indicating that abundance alone cannot explain the observed interaction patterns. Our results show that seasonal variation and temporal resolution differentially shape the importance of species attributes, highlighting the need for multi-variable approaches that account for temporal dynamics to accurately explain and predict ecological interactions.

## Introduction

Flowering plants and pollinators form complex interaction networks that are fundamental to the architecture of biodiversity (Bascompte & Jordano, 2007; Guimaraes, 2020). Yet these networks are inherently dynamic, with their structure and composition varying across space and time, limiting our ability to understand and predict species interactions (Burkle & Alarcón, 2011; Tylianakis & Morris, 2017). Documenting interactions, however, remains extremely labour-intensive (Chacoff et al., 2012), and the resulting network structures are highly dependent on sampling effort and methodology (Jordano, 2016). Necessarily then, ecological networks are often treated as static entities, with interactions aggregated across time to increase sample size and statistical power (CaraDonna et al., 2017; Poisot et al., 2015). However, the rapid turnover of species and interactions across the season means that aggregating interactions can obscure rather than clarify the processes structuring ecological systems (Duchenne et al., 2026; Payrató-Borràs et al., 2024). As a result, identifying which species attributes shape interaction patterns remains challenging because their relative importance varies across environmental conditions and time.

Improving our ability to predict species interactions is a central problem in ecology (Peralta et al., 2024; Valdovinos, 2019), particularly in the context of climate change, ongoing species loss, and the need to sustain ecosystem services. However, although overall network structure can often be partially reproduced (Crea et al., 2016; Eklöf et al., 2013), accurately predicting which species interact and the strength of their interactions remains elusive. Recent efforts to improve predictive performance have attempted to overcome limitations of conventional analytical methods, such as their inability to account for the non-independence of observations and nonlinear relationships among key variables, by incorporating species attributes such as abundance, traits and phenology (Bartomeus et al., 2016; Benadi et al., 2022). While flexible machine learning approaches have emerged as promising tools for predicting ecological interactions (Pichler et al., 2020; Strydom et al., 2021), skepticism remains regarding their general applicability, highlighting the need for comparative evaluations across ecological contexts to better understand how species attributes shape interaction patterns.

Among the species attributes proposed to shape interaction patterns, phenology, abundance, and traits have received the strongest empirical support (Peralta, Vázquez, et al., 2020; Vázquez et al., 2009). Phenology constrains the pool of potential interactions through temporal overlap (Benadi et al., 2014; Encinas-Viso et al., 2012), with approximately one-third of all potential interactions estimated to be impossible due to phenological mismatches (Olesen et al., 2011). When two species overlap in time, the probability that they interact depends on their relative abundances and traits (Olito & Fox, 2015; Stang et al., 2007), along with other abiotic (e.g. temperature) and biotic factors (e.g. interspecific competition). While abundance influences both species’ roles (Fort et al., 2016; Simmons et al., 2019) and overall network structure (Canard et al., 2014; Vázquez et al., 2009), independent estimates are often unavailable and instead inferred from interaction data (Blüthgen & Staab, 2024), potentially obscuring its contribution to interaction patterns. Likewise, morphological traits constrain the range of possible partners, and interaction probability often peaks when the interacting traits of plants and pollinators are closely matched (Klumpers et al., 2019). Yet interactions can occur despite imperfect trait matching, and multiple traits can jointly shape interaction patterns (Eklöf et al., 2013; Lanuza et al., 2023). Thus, accurately predicting plant-pollinator interactions requires integrating multiple species attributes. However, how their relative importance varies through time remains poorly understood (Ballarin et al., 2026; Barreto et al., 2025).

Most attempts to predict species interactions rely on temporally aggregated networks, potentially contributing to their limited predictive performance by obscuring temporal variation in species attributes, whose effects on interactions may become detectable only at finer temporal resolutions. In addition, species attributes may interact, generating non-additive effects on species interactions. Here, we evaluate how abundance, trait matching, and phenology shape plant-pollinator interactions and their predictability across seasonal variation and temporal resolutions. To do so, we combine complementary approaches to evaluate both the temporal and overall importance of species attributes for predicting species interactions, assess how temporal resolution influences the ability of species attributes to explain interaction network structure, and test whether pollinators exhibit preferences for particular flowering taxa beyond those expected from phenology and abundance (**Figure 1**). We collected data on plant-pollinator interactions and associated species attributes in three nearby botanical gardens with similar climatic conditions, minimizing large-scale environmental variation. This setting provides a high diversity of co-flowering species and multiple potential pairwise interactions while allowing detailed measurements of species attributes under semi-controlled conditions, providing an ideal system to test how seasonal variation and temporal resolution influence our ability to explain and predict species interactions.

**Figure 1.**
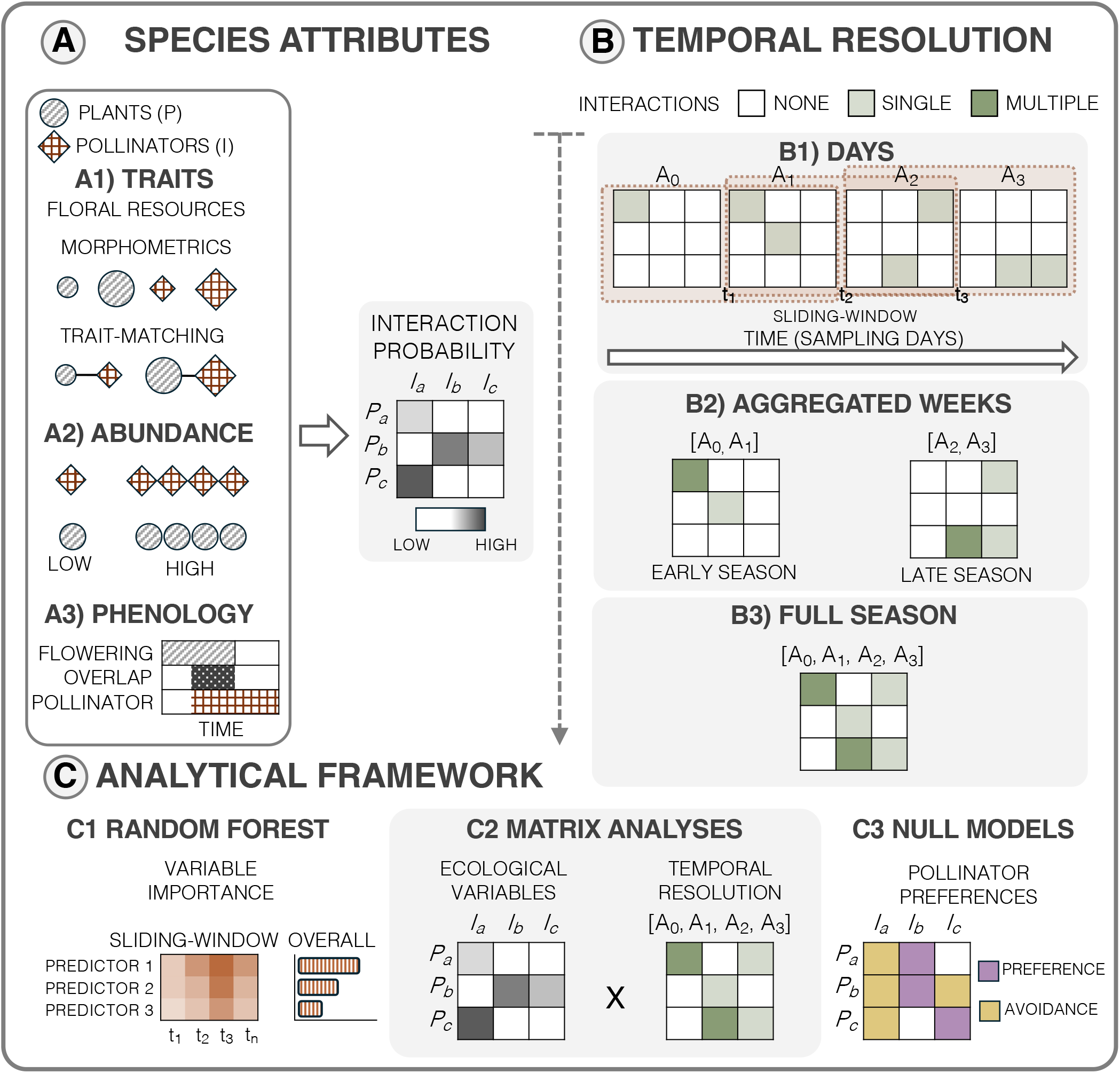
Workflow of the study showing how species attributes (A) are evaluated across temporal resolutions (B) using complementary analytical approaches (C). Panel A illustrates the species attributes examined to explain plant-pollinator interaction patterns: (A1) pollinator and floral traits, (A2) pollinator and floral abundances, and (A3) pollinator and floral phenology. Panel B presents the temporal resolution considered, including both continuous and aggregated representations: (B1) individual sampling days, treated as a continuous temporal variable, (B2) early, mid, and late season (i.e. aggregated weeks), and (B3) the full flowering season. Panel C depicts the main analytical approaches used to (C1) quantify the temporal variation and overall contribution of ecological variables to predicting interactions; (C2) assess how ecological variables and temporal resolution influence interaction network structure; and (C3) test for non-random pollinator preferences for flowering taxa using null models.

## Methods

### Study site and sampling method

We monitored plant-pollinator interactions during the entire flowering season of 2023 (23 March to 23 August) in the botanical gardens of Halle, Jena, and Leipzig (Germany; **Figure S1**). These gardens belong to the PhenObs initiative (Nordt et al., 2021), a network of botanical gardens across the Northern Hemisphere that records phenological observations on a wide range of herbaceous species. Following the PhenObs protocol, we assessed flowering species weekly and selected those that were in bloom and attracted floral visitors for focal pollinator monitoring, resulting in a set of 78 herbaceous species (hereafter focal plants; **Table S1**).

We visited each garden weekly (19-22 surveys per garden; see **Table S2**), and sampled plant-pollinator interactions during daylight hours (e.g. 09:00-19:00) for 15 minutes per focal plant under suitable weather conditions (i.e. no rain and low to moderate wind). We conducted focal observations on the same group of individual plants of each species at fixed locations within each garden from the start to the end of flowering. On each sampling day, we surveyed focal plant species in random order and recorded only pollinator individuals contacting the reproductive organs of the plant as interacting species. We use the terms “pollinator” and “floral visitor” interchangeably; although not all visitors are effective pollinators, visitation frequency provides a proxy for their relative contribution to pollination (Ballantyne et al., 2015; Vázquez et al., 2012). Pollinator monitoring began with the onset of flowering of focal plants and ended when flowering had largely ceased among focal plants in each garden, corresponding to the late phase of the season, despite continued pollinator activity and flowering of other plant species in the gardens (**Figure S2**).

To complement focal observations and capture pollinator species that may not have been recorded on focal plants, we opportunistically selected ten flowering plant species on each sampling day and observed pollinator visits to each species for 3 min (30 min per garden per sampling day). These surveys targeted non-focal plant species and focused on flowering individuals where pollinator activity was observed.

### Sampling effort, taxonomic coverage and sampling completeness

The 78 focal plant species encompassed 68 genera, 33 families, and 22 orders (**Table S1**). We conducted pollinator observations on 46 focal species in Halle (52 h), 43 in Jena (40 h), and 47 in Leipzig (49 h), with only 12 species shared among all gardens (**Table S3**). Complementary sampling of non-focal plants expanded plant coverage to 311 additional species across all gardens (i.e. unique species pooled across sites), representing 207 genera, 54 families, and 25 orders (**Table S4**). We sampled 124, 117, and 117 non-focal plant species in Halle, Jena, and Leipzig, respectively (9, 10, and 10 h per garden; **Table S5**).

In total, 243 species of floral visitors were recorded across focal and non-focal observations (168, 125, and 131 in Halle, Jena, and Leipzig, respectively), spanning 126 genera, 43 families, and 5 orders (**Table S6**). The pollinator assemblage was dominated by the main flower-visiting groups, particularly Hymenoptera (158 species) and Diptera (78), followed by Coleoptera (26), Lepidoptera (18), and Hemiptera (6). Focal observations captured 175 pollinator species across the three gardens, while non-focal observations further enriched pollinator diversity, documenting an additional 68 species not observed on focal plants. All subsequent analyses were conducted at the species level.

In total, we recorded 6,541 plant-pollinator interactions at the species level, with 2,774 in Halle, 2,132 in Leipzig, and 1,635 in Jena (including 1,262, 821, and 739 focal interactions, respectively). Most interactions involved Hymenoptera (78%) and Diptera (17%), with honeybees alone accounting for 21% of all interactions. Excluding honeybees, most interactions still involved Hymenoptera (71%) and Diptera (22%). These patterns were consistent across the three gardens. Overall, the two sampling approaches provided a comprehensive overview of the pollinator assemblages within these botanical gardens. Accumulation curves (**Figure S3**) indicate that sampling effort was sufficient to characterize pollinator assemblages associated with focal plants across gardens. However, observations of non-focal plants proved more efficient at capturing overall pollinator diversity at the garden scale, yielding a more complete representation of the pollinator assemblage with lower sampling effort while covering a broader range of plant species.

### Data description

To evaluate the different mechanisms that drive plant-pollinator interactions, detailed information on abundance, phenology and species traits for both plants and pollinators was recorded as follows:

i. During each sampling event in each botanical garden, we recorded floral abundance for each sampled plant (from both focal and non-focal observations) by counting the total number of open flowers or inflorescences on the sampled individuals selected for pollinator observations, depending on floral morphology. Species with dense inflorescences composed of numerous small florets (primarily Asteraceae and Caprifoliaceae) were assessed at the inflorescence rather than flower level (6% of focal species; 15% of all flowering species considered). Flower size was measured separately to account for variation in flower size across species (see below). We also recorded the number of pollinator individuals visiting each sampled plant during observation periods. Pollinator individuals were distinguished in the field as morphospecies during focal observations. To minimise collecting while ensuring reliable taxonomic identification, we captured a single representative individual of each morphospecies from each sampled plant species, collecting additional individuals only when required to confirm identifications. This approach allowed us to quantify both the number of pollinator individuals and their visitation frequency to each sampled plant. Although repeated visits by the same individual cannot always be distinguished, particularly under high visitation rates, this approach provides a closer approximation of pollinator abundance than inferring abundance from visitation frequencies alone (Blüthgen et al., 2006; Vizentin-Bugoni et al., 2014).
ii. Plant phenology was obtained from the PhenObs project, which provides weekly records of flowering intensity for all focal species (Nordt et al., 2021). In contrast, pollinator phenology cannot be directly inferred from interaction data alone, as observations are inherently stochastic and may not capture the full flight period of each species. To address this, we complemented our observational data with occurrence records from the Global Biodiversity Information Facility (GBIF). Germany has high coverage of occurrence data, particularly in urban areas such as botanical gardens (Rocha-Ortega et al., 2021; Sweet et al., 2022). We extracted occurrence records for each pollinator species from the same sampling year within a region centred on the three study cities, defined by a rectangular extent of approximately 1000 km (east-west) and 500 km (north-south). This spatial extent represented a compromise between maximizing sample size and limiting climatic heterogeneity (**Figure S4**). This resulted in 2,933,772 occurrence records (https://doi.org/10.15468/dl.k5cnuf). Pollinator phenology was thus standardised across gardens; although this approach has limitations, it provides a more robust estimate than relying solely on interaction data and closely matches the temporal distribution of observed interactions (e.g. **Figure S5**).

Plant and pollinator phenology curves were estimated using species-specific generalized additive models (GAMs). Plant models were fitted to weekly flowering records from the PhenObs project, whereas pollinator models were fitted to daily GBIF occurrence records supplemented with our own pollinator observations. Pollinator activity periods were defined as the interval between the first and last days on which predicted relative activity exceeded 10% of the maximum predicted activity for each species. Plant and pollinator phenology curves were then used to construct phenology-based interaction probability matrices. In addition, the number of days during which the plant flowering period and the pollinator activity period overlapped (i.e. phenological overlap) was calculated and used as a predictor of interaction frequency in the random forest and GAM analyses.

1. We measured plant traits associated with plant-pollinator interactions and reproductive strategies (E-Vojtkó et al., 2020; Lanuza et al., 2023), including flower number, plant height, style length, flower size, ovule number, autonomous selfing, flower lifespan, flowering duration, nectar volume, and pollen production per flower (**Table S7a**). Most traits were measured using three replicates per species, while nectar volume and pollen production were measured using five replicates per species. Nectar volume was quantified in freshly opened flowers that were bagged prior to anthesis. Autonomous selfing was quantified from a variable number of observations per species (mean = 19). Because it was the only trait not measured for all species, literature data were used to supplement estimates of selfing for 47

We measured pollinator morphological traits including intertegular distance (for bees), body width (for non-bee taxa), body length, and proboscis length (**Table S7b**), which are key morphological traits mediating plant-pollinator interactions (Földesi et al., 2021; Klumpers et al., 2019). We measured intertegular distance (ITD), body width and body length from images using ImageJ. We estimated proboscis length for bee species using the pollimetry package (Kendall et al., 2019), and for non-bee species using values from the literature and, when possible, from images. Intertegular distance in bees and body width in non-bee taxa were treated as comparable estimates of pollinator body size in subsequent analyses. Although body length and proboscis length were also measured, trait matching analyses focused exclusively on pollinator body size because it was directly measured for all taxa and was highly correlated with both body length and proboscis length (r = 0.76-0.77), reducing redundancy among morphological traits.

Trait matching between plants and pollinators was quantified using a Gaussian matching function based on pollinator body size and flower width 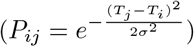, where *P*_*ij*_ represents the interaction probability between plant *i* and pollinator *j, T*_*i*_ and *T*_*j*_ are their respective trait values, and *σ* is a tolerance parameter controlling the decline in interaction probability with increasing morphological mismatch (Bartomeus et al., 2016; Williams et al., 2010).

### Seasonal phenological patterns

To characterize seasonal phenological patterns within the botanical gardens, we examined species-level phenology and community-level flowering synchrony. We first examined the phenological distribution of plant and pollinator species based on their phenological centres, defined as the weighted mean day of year (DOY), with weights corresponding to flowering intensity (plants) or activity probability (pollinators). Species with fewer than three observations were excluded. Because pollinator phenology was estimated at the regional level rather than separately for each garden, a single phenological distribution was used across all gardens. To assess whether flowering phenology differed across botanical gardens, we compared the flowering centroids of plant species shared across gardens using a linear mixed-effects model, with garden as a fixed effect and species identity as a random effect. In addition, for each garden, we quantified community flowering synchrony using a simplified proxy of synchrony based on the temporal spread of species phenological centres, calculated as 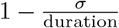, where duration was defined as the difference between the 95th and 5th percentiles of species phenological centres. We also quantified the standard deviation (*σ*) of phenological centres within each garden. Higher synchrony values indicate stronger temporal clustering among species.

### Analyses

The effects of abundance, phenology, and trait matching on plant-pollinator interactions were evaluated across the flowering season and at multiple temporal resolutions (i.e. sampling-day, weekly aggregated, and full-season networks) using complementary analytical approaches. First, we quantified how the importance and predictive performance of species attributes shaping interactions varied through the season and across the full flowering period (C1, **Figure 1**). We also modelled visitation dynamics to estimate the direction and magnitude of ecological variable effects on plant-pollinator interactions. Next, we examined how variable-specific probability matrices and temporal resolution influenced interaction network structure (C2). Finally, we used null models accounting for spatiotemporal variation and abundance to test whether pollinators exhibited non-random preferences for particular plant taxa, potentially reflecting the influence of unmeasured traits (C3). To complement this analysis, we quantified species-level selectivity to assess how pollinator preferences change through time, comparing estimates based on interaction frequencies alone with those accounting for in situ floral abundances. For all analyses, focal and non-focal observations of the same plant species within the same garden and sampling day were jointly considered when both observation types were available; otherwise, only focal observations were used.

#### Temporal and overall importance of species attributes

We used random forest models to assess the relative importance of ecological variables and their ability to predict plant-pollinator visitation rates. Random forest was used to capture complex non-linear relationships among predictors while providing robust estimates of variable importance (Breiman, 2001). Two complementary approaches were implemented: a sliding-window approach to evaluate how the importance of species attributes for visitation patterns changes across the season using sampling day as a continuous temporal variable, and a global model to estimate the overall importance of species attributes.

Models were fitted using the ranger package (Wright et al., 2019) on weekly interaction data pooled across the three botanical gardens, with visitation rate modelled as log(1 + Visitation rate). Preliminary analyses indicated that garden identity explained negligible variation in interaction frequency and was therefore not included as a predictor. Predictor variables included floral abundance, pollinator abundance, flower width, trait matching, and phenological overlap, quantified as the number of days during which the flowering period of a plant overlapped with the activity period of a pollinator species. Random forests were fitted with 1,500 trees, and predictor importance was quantified using SHAP (SHapley Additive exPlanations) values computed with the fastshap package (Greenwell & Greenwell, 2020) and summarised as mean absolute SHAP values across observations. Predictive performance was evaluated using an 80/20 stratified train-test split and quantified using the coefficient of determination (R2), root mean square error (RMSE), and mean absolute error (MAE). Temporal variation in variable importance was assessed using a sliding-window approach. Models were fitted within moving windows of five consecutive weeks (step = 1 week), using the same predictor variables as in the global model. Sampling week was excluded as a predictor to avoid confounding temporal trends with variable effects, and windows containing fewer than 80 observations were excluded to ensure model stability. Within each window, predictor importance was quantified using SHAP values and standardised to obtain relative importance, allowing comparison of variable contributions through time. The global model was fitted using the complete seasonal dataset and was used to estimate the overall relative importance of species attributes across the flowering season. To complement the random forest analyses, we fitted generalized additive mixed models (GAMMs) using the mgcv package (Wood & Wood, 2015). The same predictor variables as in the random forest analyses were included to estimate the direction and statistical support of predictor effects, while testing interactions between pollinator abundance and both floral abundance and floral width.

#### Variation in species-attribute effects across temporal resolution

To evaluate how ecological variables influence visitation patterns across temporal resolutions, interaction and variable-specific matrices were constructed at three temporal levels for each botanical garden (sampling days, aggregated weeks, and the full seasonal network). Visitation-rate networks were built for pollinators identified to species level and for the main pollinator orders observed in the gardens (Hymenoptera, Diptera, Coleoptera, Lepidoptera). Plant-pollinator interaction records were aggregated by garden, sampling day, and species pair to generate weighted bipartite interaction matrices. Visitation rate was first calculated at the sampling-day level by dividing total interactions for each plant-pollinator pair by the total observation time for each plant species within a garden. These standardized rates were subsequently summed across sampling occasions, when required for weekly and full-season aggregations, to produce garden-level visitation-rate networks representing cumulative interaction intensity while accounting for plant-specific sampling effort.

Abundance probability matrices were generated for each botanical garden and temporal resolution using standardized relative abundances of pollinator and flowering species. Pollinator and floral abundances were summed within each garden and temporal unit and converted to relative abundances by dividing species-level abundances by total community abundance. Interaction probabilities were calculated by combining plant and pollinator relative abundances under an independence assumption, with expected interaction probabilities proportional to the product of plant and pollinator relative abundances (Vázquez et al., 2009).

Unlike the phenological-overlap predictor used in the random forest and GAMM analyses, which was based on the overlap in activity periods, phenology probability matrices were derived from the same species-specific temporal activity distributions but represented the probability of temporal co-occurrence. Plant and pollinator probabilities were then combined under an independence assumption by multiplying their respective probabilities for each sampling week. Resulting interaction probabilities for each plant-pollinator pair were subsequently averaged for aggregated temporal resolutions (i.e. aggregated weeks or full season networks).

Trait-based probability matrices were constructed using the Gaussian trait matching function described above. A global plant × pollinator trait matching matrix was first assembled using all plant and pollinator species and then filtered to include only the species present in each botanical garden and temporal resolution.

We used matrix similarity analyses to assess how closely probability matrices derived from ecological variables reproduced observed visitation matrices across temporal resolutions within each botanical garden. Results were summarized across gardens to identify general temporal patterns. We assessed matrix similarity using Procrustes analysis implemented through the protest function in the vegan package (Oksanen et al., 2007), and the strength of configuration similarity was quantified using the Procrustes correlation coefficient (r). As a robustness check, analogous analyses were conducted using Mantel tests. Because probability matrices for all variables were scaled between 0 and 1, Euclidean distance was used as the distance metric, as it preserves the geometric relationships among observations.

#### Pollinator preferences

To identify interaction patterns that could not be explained by plant and pollinator abundances and sampling structure alone, we compared observed interaction frequencies with expectations derived from null models. To improve statistical power and facilitate the interpretation of broad interaction patterns, interaction data were aggregated at the level of plant families and pollinator guilds. Interaction matrices were then constructed separately for each botanical garden and sampling week to preserve the spatiotemporal structure of the data. Plant families and pollinator guilds represented by fewer than 50 total interactions across the dataset were excluded to ensure robust estimation. Within each garden-week matrix, interactions were randomized using the Patefield’s algorithm, implemented in r2dtable function in the bipartite package (Dormann et al., 2008), which maintains row and column totals and thus preserves the total number of interactions per plant family and pollinator guild while randomizing partner identities. For each matrix, 1,000 randomized networks were generated, and randomized matrices were subsequently summed across garden-week units to obtain expected interaction frequencies for each plant family-pollinator guild pair under spatiotemporal constraints. As a sensitivity analysis, we repeated the null-model procedure using alternative minimum interaction thresholds (25 and 75 interactions) and confirmed that the main preference and avoidance patterns remained qualitatively consistent.

Deviations from null expectations were quantified using standardized effect sizes (SES), calculated as the difference between observed and expected interaction frequencies divided by the standard deviation of the null distribution. Positive SES values indicate associations (more interactions than expected), whereas negative values indicate avoidance. Statistical significance was assessed using two-tailed p-values derived from the simulated distributions, applying a conservative threshold of p < 0.01. Interactions that were never observed to co-occur within the same garden and sampling week were classified as spatiotemporally impossible. We also conducted analogous analyses at a finer taxonomic resolution for pollinators, using pollinator families and genera. Finally, we tested whether trait matching probability was associated with deviations from null expectations at the plant family × pollinator genus level as a complementary analysis.

To better understand temporal variation in pollinator preferences, we calculated weekly species-level selectivity (*d*^*′*^; Blüthgen et al., 2006) from interaction frequency matrices using the bipartite package. Because *d*^*′*^ measures deviation from random interactions given partner availability, we additionally calculated an abundance-weighted version of *d*^*′*^ by standardizing interaction frequencies by floral abundance to assess how resource availability influences seasonal patterns of specialization.

#### Software and use of large language models

All analyses were conducted in R v4.5.0 (R Core Team, 2025). ChatGPT (OpenAI) was used during manuscript preparation to assist with code development, debugging, and language refinement. All outputs were reviewed and validated by the authors, who are solely responsible for the ideas, analyses, interpretations, and conclusions presented in this manuscript.

## Results

### Seasonal phenological patterns

Flowering phenology was similar across botanical gardens, with mid-season phenological centres occurring in mid season (DOY range = 145-157), comparable flowering durations among sites (range = 107-116 days; **Figure S7**), and moderate synchronization across gardens (range = 0.63-0.68). This consistency was further supported by a species-level comparison of flowering centroids for species shared across gardens, which revealed a negligible effect of garden on flowering timing (F = 0.06), while species identity explained 89% of the variation in flowering time. Pollinator phenology, derived from regional GBIF records, showed a similar degree of temporal clustering (synchronization = 0.64) but peaked later in the season than flowering phenology (DOY = 181), despite a comparable phenological spread (duration = 100 days; **Figure S7**).

### Temporal and overall importance of species attributes

The overall random forest model displayed moderate to high predictive performance (R2 = 0.65, RMSE = 0.66, MAE = 0.44), indicating that the selected predictors captured a substantial proportion of the variation in visitation rates. Variable importance differed markedly among predictors. Pollinator abundance showed the highest overall importance, followed by trait matching and floral abundance, whereas floral width had a more moderate contribution. Phenological overlap and nectar volume contributed the least overall (**Figure 2b**).

**Figure 2.**
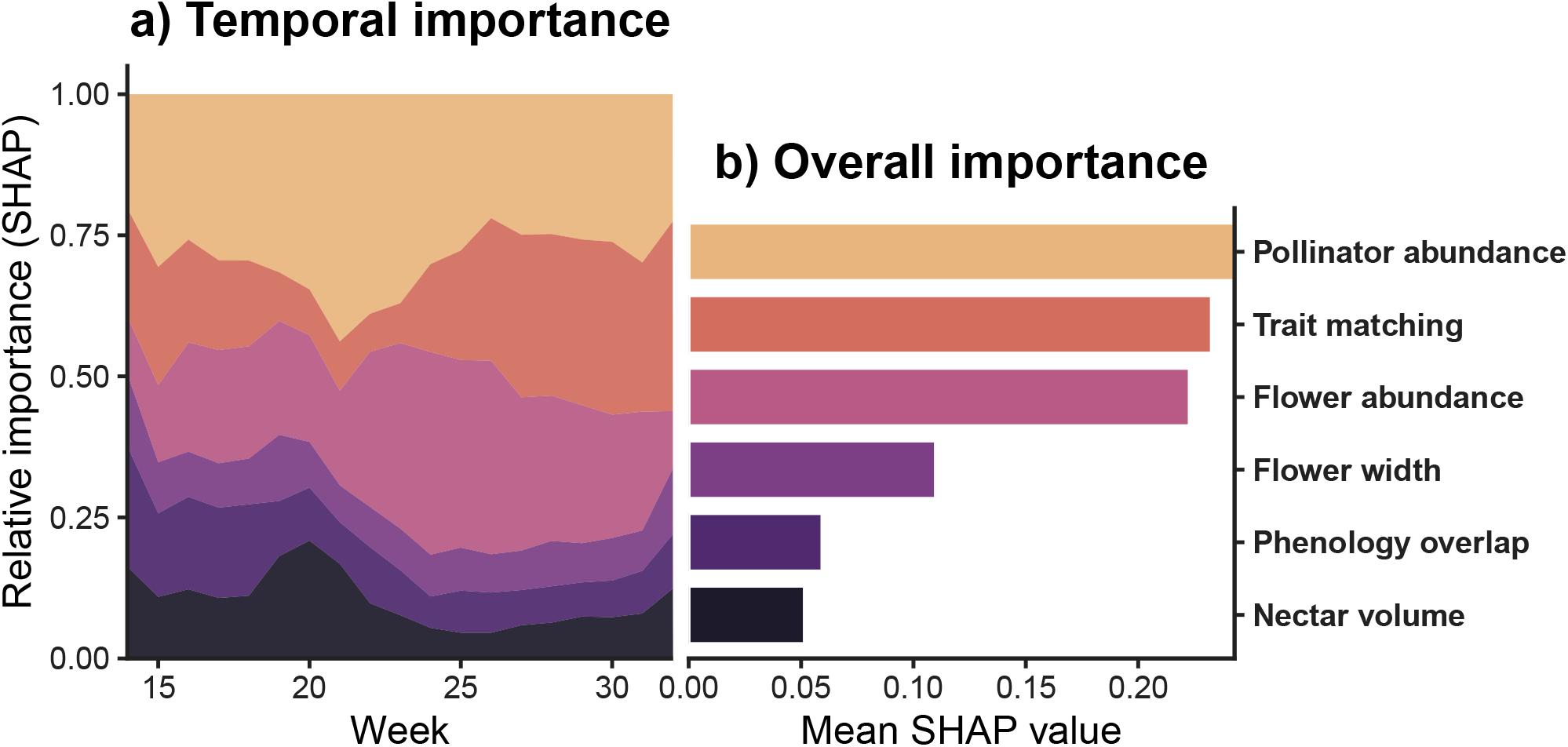
Temporal and overall importance of species attributes for predicting pollinator visitation rates. Relative importance of species attributes estimated from random forest models using SHAP (SHapley Additive exPlanations) values. (a) Weekly variation in the relative contribution of each predictor to model predictions across the flowering season. Values were estimated using sliding-window random forest models fitted over 5-week intervals and assigned to the central week of each window. (b) Overall importance of each predictor summarized as the mean SHAP value across all weeks.

The relative importance of ecological variables also changed across the season (**Figure 2a**). Pollinator abundance was consistently among the most important predictors and peaked during periods of highest activity. In contrast, trait matching showed substantial seasonal variation, tending to gain importance when abundance-based predictors were less dominant. Floral abundance also exhibited pronounced variation, showing the largest temporal variation among predictors while contributing substantially across the season. Floral width had a more modest but relatively consistent contribution through time, whereas phenological overlap and nectar volume remained comparatively low throughout the season. Overall, the GAMM results were consistent with the random forest analyses, confirming the dominant roles of pollinator and floral abundance, while revealing a weaker positive effect of trait matching and interactions between pollinator abundance and both floral abundance and flower width (**Figures S8b-d**).

### Variation in species-attribute effects across temporal resolution

Across temporal aggregation levels, abundance-based probability matrices consistently showed the strongest similarity to observed interaction frequency matrices (**Figure 3**), with Procrustes correlations increasing from moderate values at weekly resolution (r = 0.6, range = 0.4-0.85) to high values at the full-season resolution (r = 0.84, range = 0.67-0.94) across all botanical gardens. In addition, within-season patterns revealed that the influence of abundance strengthened towards the later stages of the season. Trait matching matrices exhibited intermediate similarity, with relatively consistent correlations across temporal resolutions (range = 0.29-0.8), but tended to peak at intermediate levels of temporal aggregation (i.e. aggregated weeks, r = 0.7, range = 0.61-0.8). In contrast, phenology-based matrices showed moderate similarity at finer temporal resolution (r = 0.51, range = 0.43-0.6) but did not increase similarity with temporal aggregation and tended to weaken at the full-season level (r = 0.21). These temporal patterns were qualitatively consistent across study sites (**Figure S9**), indicating that the contribution of species attributes to interaction structure varies with temporal resolution. Analogous analyses based on Mantel tests yielded similar results (**Figure S10**).

**Figure 3.**
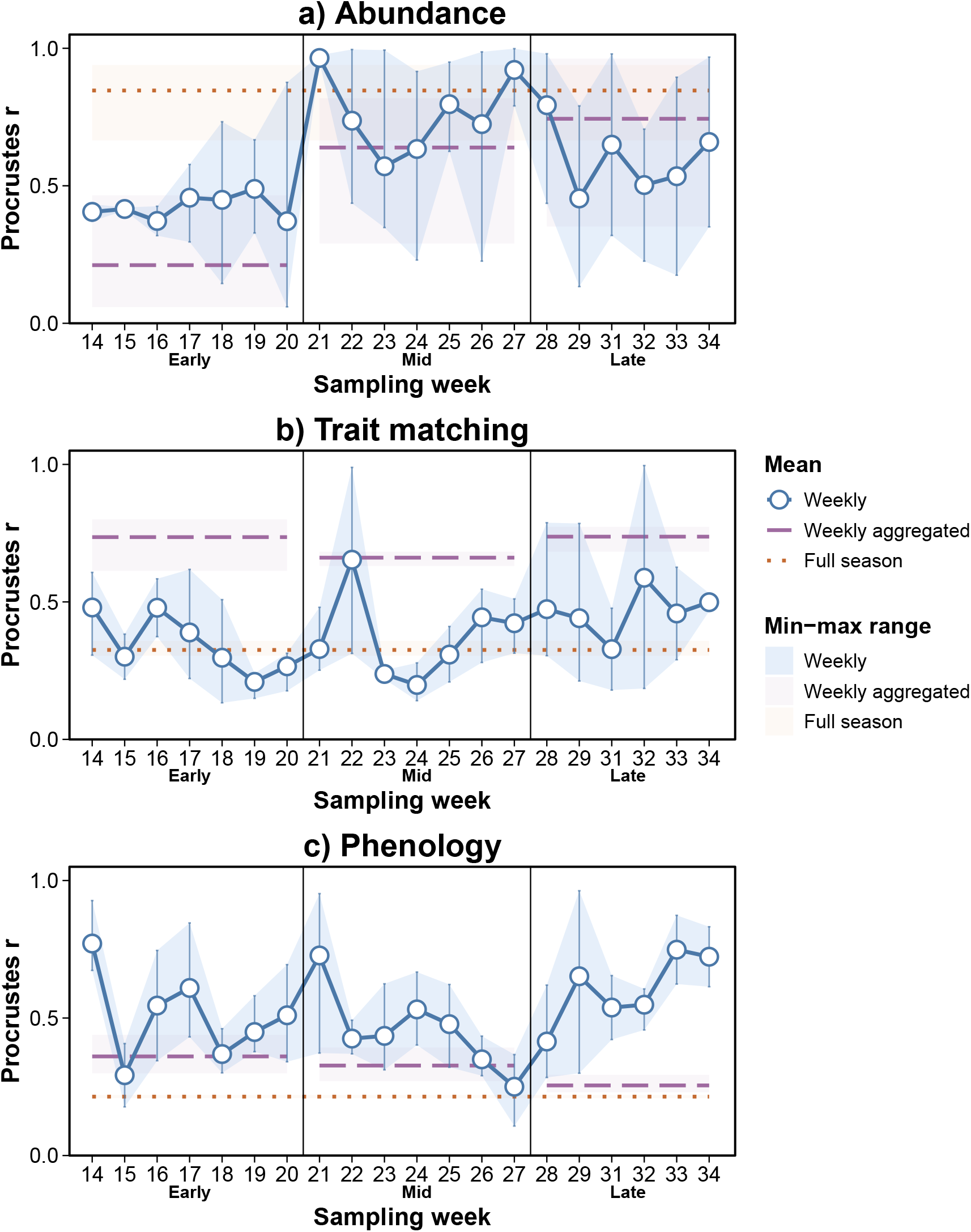
Procrustes association between visitation rate and plant-pollinator interaction probability matrices derived from abundance (a), trait matching (b), and phenology (c) across levels of temporal resolution (weekly, weekly aggregated, and full season), while accounting for variability among botanical gardens. Weekly aggregated values correspond to three seasonal categories (early, mid, and late), obtained by pooling weekly networks into three equal temporal partitions of the full sampling period. Weekly values represent individual weekly networks within each seasonal category. Solid lines and points represent mean Procrustes associations for weekly networks across botanical gardens, whereas dashed and dotted horizontal lines indicate weekly aggregated seasonal networks and full-season networks, respectively. Shaded areas and error bars indicate the minimum-maximum range among botanical gardens.

### Pollinator preferences

Null model analyses revealed non-random patterns of pollinator preference and avoidance among pollinator genera and plant families (**Figure 4a**). Overall, 58.6% of interactions did not deviate from null expectations, whereas 15.7% showed positive associations (preference) and 25.7% exhibited negative associations (avoidance; p < 0.01). A substantial proportion of potential interactions (25.4%) were spatiotemporally impossible, highlighting the strong constraints imposed by species turnover through space and time. Preference patterns varied markedly among pollinator guilds, with some exhibiting consistent associations with a limited set of plant families, whereas others showed broader and more diffuse interaction patterns (**Figure 4b**). Despite this variation, most pollinator guilds interacted with multiple plant families, consistent with broadly generalist interaction patterns shaped by non-random preferences. These patterns were qualitatively consistent across alternative pollinator taxonomic resolutions (**Figure S11** and **Figure S12**). Trait matching probability showed a weak positive association with the magnitude of deviations from null expectations (**Figure S13**).

**Figure 4.**
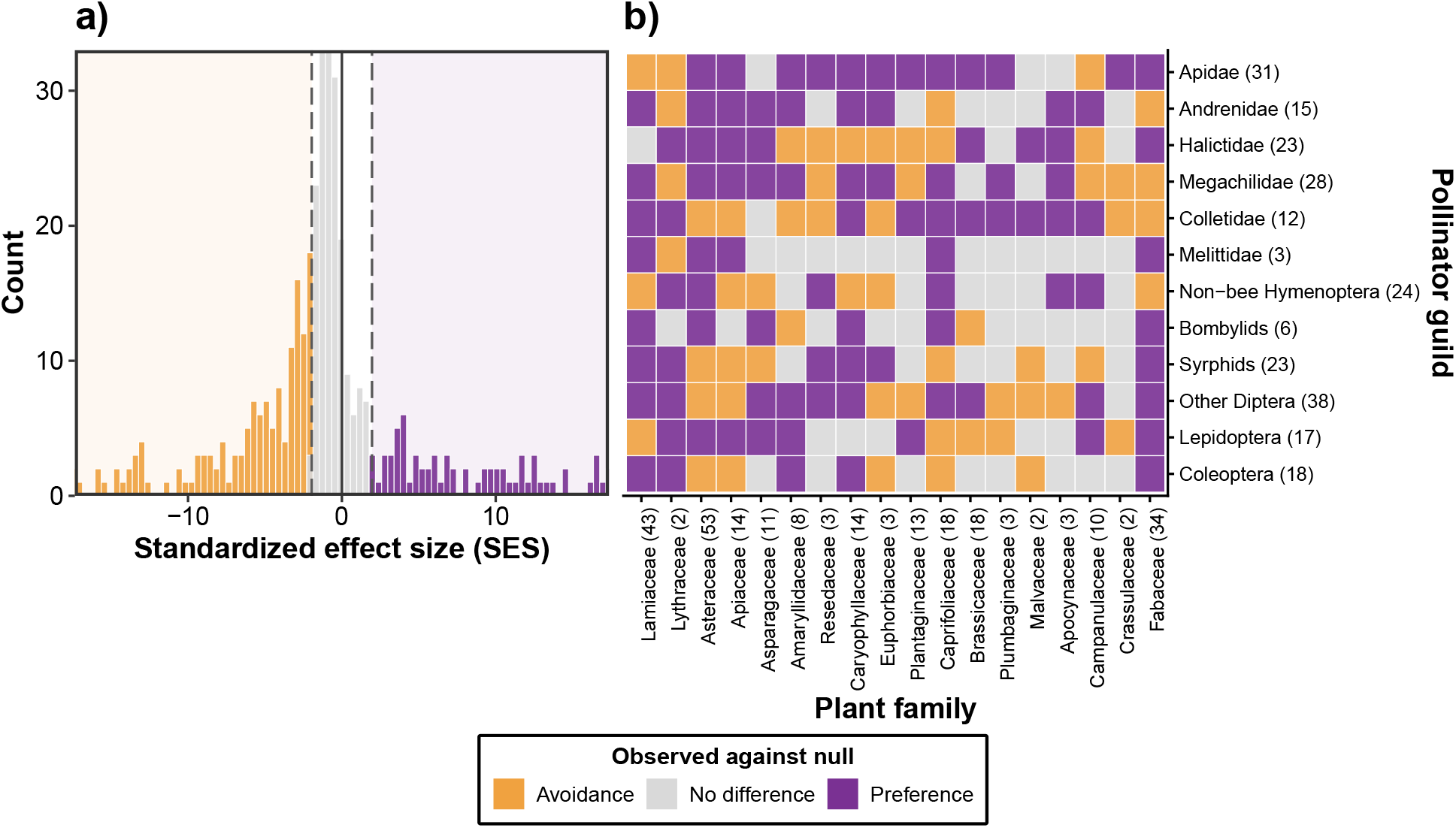
Standardized effect sizes (SES) of plant-pollinator interactions relative to null expectations derived from week- and garden-specific randomizations. Colours denote deviation from null expectations (avoidance, no difference, preference). a, Distribution of SES values across all plant-pollinator genus pairs; dashed lines indicate critical values (±1.96) b, Interaction patterns at the plant family × pollinator guild level for the 18 plant families with the strongest overall SES signal (sum of absolute SES values). Numbers in parentheses indicate species richness for each pollinator guild (y axis) and plant family (x axis).

Pollinator specialization (*d*^*′*^) showed substantial temporal variation across botanical gardens (**Figure S14**). However, abundance-weighted *d*^*′*^ values showed lower overall selectivity and reduced seasonal variation, indicating that floral abundance strongly shapes apparent specialization patterns.

## Discussion

Our results show that the influence of species attributes on plant-pollinator interactions is not constant through time, supporting the view that interaction structure emerges from temporally dynamic processes (Ballarin et al., 2026; Barreto et al., 2025). Pollinator abundance, floral abundance, and trait matching consistently emerged as the most influential predictors of plant-pollinator interactions across the season. Abundance-driven processes dominated during most of the season, peaking around the phenological centres of plants and pollinators, whereas trait matching gained relative importance as the influence of abundance decreased, primarily refining visitation rates rather than overall network structure. Importantly, a relatively small set of species attributes captured much of the variation in visitation patterns, as reflected by the moderate to high predictive performance of the models, supporting the view that complex ecological dynamics can be predicted from a limited set of key variables (Peralta et al., 2024). In addition, persistent patterns of preference and avoidance across taxonomic levels indicate that interactions cannot be fully explained by spatiotemporal overlap and abundance alone. Overall, these findings suggest that interaction dynamics can be understood through a combination of dominant processes and context-dependent constraints, supporting a general framework in which abundance governs interaction frequencies, while traits and phenology constrain which interactions are feasible, with their relative influence varying across temporal contexts.

These findings reinforce previous evidence that species abundances are major determinants of plant-pollinator interaction structure (Dormann et al., 2017; Fort et al., 2016; He et al., 2025; Simmons et al., 2019). However, this effect is strongly context-dependent, as floral abundance translates into higher visitation only when pollinators are abundant, with pollinator generalisation increasing during periods of peak phenological overlap. Seasonal variation in mean species selectivity indicates that interaction roles emerge from changing community composition and species abundances rather than representing fixed species properties (Brosi, 2016). In addition, when floral abundance is explicitly accounted for, apparent specialisation is reduced, indicating that much of the observed selectivity reflects variation in floral abundance rather than shifts in interaction preferences. This highlights the need for caution when inferring interaction patterns in the absence of independent abundance data, a limitation that remains common in ecological network studies (Blüthgen & Staab, 2024; Olito & Fox, 2015). However, consistent non-random interaction patterns indicate that additional ecological processes, although comparatively weaker than the effects of species abundance, also contribute to structuring plant-pollinator interactions (Gómez-Martínez et al., 2022; Kelly & Elle, 2021; Morán-López et al., 2022).

Trait matching exerted a consistent, intermediate influence on interactions across the season, suggesting a temporally stable constraint on interactions. While abundance shapes overall interaction patterns, trait compatibility imposes persistent constraints on which interactions can occur (Bartomeus et al., 2016; Kaiser-Bunbury et al., 2014). This is consistent with evidence across systems, including the correspondence between pollinator morphology and floral traits, such as the matching between proboscis length and floral tube depth in Hymenoptera (Klumpers et al., 2019; Zhao et al., 2022), as well as broader macroecological patterns linking hummingbird bills and floral morphology across biomes (Sánchez-Martín et al., 2025). In our study, size matching (i.e. body size-flower size) was used as a proxy for trait compatibility, as key traits such as tongue length were unavailable for many taxa and often required extrapolation. This aligns with previous work highlighting body size as a strong predictor of interactions, while also suggesting that predictive accuracy improves when multiple traits are considered jointly (Eklöf et al., 2013). Interestingly, trait matching was assigned lower importance by GAMs than by random forests in our analyses, suggesting that more flexible modelling approaches may better capture its contribution to plant-pollinator interactions, consistent with previous findings (Pichler et al., 2020). The comparatively weaker role of floral resources is consistent with a hierarchical filtering process, in which trait compatibility constrains feasible interactions, and resource-related factors influence interaction strength only once this constraint is met (Junker et al., 2013).

Phenology played a limited role in structuring interactions in this system and weakened with temporal aggregation, despite filtering feasible interactions through temporal co-occurrence. This pattern may reflect both the loss of fine-scale temporal structure through aggregation (CaraDonna et al., 2017; Schwarz et al., 2020), and the high richness of flowering species and broad phenological overlap observed in this system, which can relax phenological constraints and increase the range of feasible interactions (Benadi et al., 2014; de Manincor et al., 2020). These dynamics may be particularly pronounced in urban or human-modified systems, where extended flowering periods can buffer seasonal declines and maintain relatively stable pollinator richness across the season (Leong et al., 2016). By contrast, in systems with stronger phenological filtering (e.g. arid or alpine ecosystems), phenology plays a stronger role in structuring interactions and fitness outcomes (Encinas-Viso et al., 2012; Vázquez et al., 2023), and it may provide stronger predictive power across seasons than within-season interaction dynamics (Bramon Mora et al., 2020; CaraDonna et al., 2021; Peralta, Perry, et al., 2020). In addition, the broader-scale estimation of pollinator phenology highlights the need for temporally and spatially resolved measurements to better capture the fine-scale dynamics shaping ecological interactions.

Although botanical gardens differ from natural habitats by increasing overlap in flowering times and the number of potential interaction combinations, they provide a valuable system for investigating plant-pollinator interactions under replicated and semi-controlled conditions across phylogenetically and functionally diverse species assemblages. At the same time, the low genetic diversity and frequent clonality of plants in botanical gardens may reduce the contribution of individual variation to interactions (Arroyo-Correa et al., 2021; Kuppler et al., 2016). Beyond within-season dynamics, interactions can also rewire across years (Domínguez-García et al., 2026; Resasco et al., 2021), highlighting the importance of incorporating interannual variability and climatic fluctuations when evaluating interaction patterns. More broadly, the processes structuring species interactions are likely to vary across environments and taxonomic groups. For example, alpine communities can be more strongly shaped by phenological dynamics (Kudo & Cooper, 2019), whereas tropical systems and specialized pollination networks such as hawkmoth-plant (Sazatornil et al., 2016) and hummingbird-plant interactions (Dalsgaard et al., 2021) can be more strongly structured by niche-based processes and trait matching (Schemske et al., 2009). Together, these patterns highlight the need for integrative and temporally explicit approaches to better understand ecological interaction dynamics. Overall, our re-sults show that plant-pollinator interactions emerge from a limited set of species attributes whose relative importance shifts through time. While abundance governs encounter rates, trait matching and phenology constrain which interactions are feasible, generating dynamic but structured interaction patterns. By integrating high-resolution temporal data, independent estimates of abundance, and complementary analytical approaches, this study demonstrates that these processes can be disentangled and quantified even in highly dynamic systems. This provides a general framework for understanding species interactions and highlights the importance of incorporating temporal dynamics and independent ecological information to better predict how ecological networks may respond to environmental change.

## Acknowledgements

We are grateful to the staff of the Botanical Gardens for their support, especially Janin Naumann (Jena), Rolf A. Engelmann (Leipzig), Martin Freiberg (Leipzig) and Norbert Schirrmeister (Halle), as well as Dennis Böttger for his assistance with fieldwork in Jena. We also thank Ignasi Bartomeus for valuable comments and suggestions. JBL was supported by a Juan de la Cierva fellowship funded by the Spanish Ministry of Science and Innovation and by the Federal State of Saxony-Anhalt through the MLU-BioDivFund program. This study was carried out within the framework of the Spanish Government’s activities through the ‘María de Maeztu Centre of Excellence’ accreditation to IMEDEA (CSIC-UIB) (CEX2021-001198).

## References

Arroyo-Correa, B., Bartomeus, I., & Jordano, P. (2021). Individual-based plant–pollinator networks are structured by phenotypic and microsite plant traits. Journal of Ecology, 109 (8), 2832–2844.

Ballantyne, G., Baldock, K. C. R., & Willmer, P. G. (2015). Constructing more informative plant–pollinator networks: Visitation and pollen deposition networks in a heathland plant community. Proceedings of the Royal Society B: Biological Sciences, 282 (1814), 20151130.

Ballarin, C. S., Amorim, F. W., Arroyo-Correa, B., & Jordano, P. (2026). Floral resource diversity drives spatiotemporal variation in plant–pollinator network structure. Oikos, 2026 (6), e11730.

Barreto, E., Duchenne, F., Beck, H., Bello, C., Bobato, R., Brenes, E., Bôlla, D., Büttner, N., Caron, A. P., Castro, J. A., et al. (2025). Hummingbird flower visitation rates vary with species traits, floral abundance and phenology across bioregions. Oikos, 2025 (9), e11354.

Bartomeus, I., Gravel, D., Tylianakis, J. M., Aizen, M. A., Dickie, I. A., & Bernard-Verdier, M. (2016). A common framework for identifying linkage rules across different types of interactions. Functional Ecology, 30 (12), 1894–1903.

Bascompte, J., & Jordano, P. (2007). Plant-animal mutualistic networks: The architecture of biodiversity. Annual Review of Ecology, Evolution, and Systematics, 38 (1), 567–593.

Benadi, G., Dormann, C. F., Fründ, J., Stephan, R., & Vázquez, D. P. (2022). Quantitative prediction of interactions in bipartite networks based on traits, abundances, and phylogeny. The American Naturalist, 199 (6), 841–854.

Benadi, G., Hovestadt, T., Poethke, H.-J., & Blüthgen, N. (2014). Specialization and phenological synchrony of plant–pollinator interactions along an altitudinal gradient. Journal of Animal Ecology, 83 (3), 639–650.

Blüthgen, N., Menzel, F., & Blüthgen, N. (2006). Measuring specialization in species interaction networks. BMC Ecology, 6, 1–12.

Blüthgen, N., & Staab, M. (2024). A critical evaluation of network approaches for studying species interactions. Annual Review of Ecology, Evolution, and Systematics, 55.

Bramon Mora, B., Shin, E., CaraDonna, P. J., & Stouffer, D. B. (2020). Untangling the seasonal dynamics of plant–pollinator communities. Nature Communications, 11 (1), 4086.

Breiman, L. (2001). Random forests. Machine Learning, 45 (1), 5–32.

Brosi, B. J. (2016). Pollinator specialization: From the individual to the community. New Phytologist, 210 (4), 1190–1194.

Burkle, L. A., & Alarcón, R. (2011). The future of plant–pollinator diversity: Understanding interaction networks across time, space, and global change. American Journal of Botany, 98 (3), 528–538.

Canard, E., Mouquet, N., Mouillot, D., Stanko, M., Miklisova, D., & Gravel, D. (2014). Empirical evaluation of neutral interactions in host–parasite networks. The American Naturalist, 183 (4), 468–479.

CaraDonna, P. J., Burkle, L. A., Schwarz, B., Resasco, J., Knight, T. M., Benadi, G., Blüthgen, N., Dormann, C. F., Fang, Q., Fründ, J., et al. (2021). Seeing through the static: The temporal dimension of plant–animal mutualistic interactions. Ecology Letters, 24 (1), 149–161.

CaraDonna, P. J., Petry, W. K., Brennan, R. M., Cunningham, J. L., Bronstein, J. L., Waser, N. M., & Sanders, N. J. (2017). Interaction rewiring and the rapid turnover of plant–pollinator networks. Ecology Letters, 20 (3), 385–394.

Chacoff, N. P., Vázquez, D. P., Lomáscolo, S. B., Stevani, E. L., Dorado, J., & Padrón, B. (2012). Evaluating sampling completeness in a desert plant–pollinator network. Journal of Animal Ecology, 81 (1), 190–200.

Crea, C., Ali, R. A., & Rader, R. (2016). A new model for ecological networks using species-level traits. Methods in Ecology and Evolution, 7 (2), 232–241.

Dalsgaard, B., Maruyama, P. K., Sonne, J., Hansen, K., Zanata, T. B., Abrahamczyk, S., et al. (2021). The influence of biogeographical and evolutionary histories on morphological trait-matching and resource specialization in mutualistic hummingbird–plant networks. Functional Ecology, 35 (5), 1120–1133.

de Manincor, N., Hautekeete, N., Piquot, Y., Schatz, B., Vanappelghem, C., & Massol, F. (2020). Does phenology explain plant–pollinator interactions at different latitudes? An assessment of its explanatory power in plant–hoverfly networks in French calcareous grasslands. Oikos, 129 (5), 753–765.

Domínguez-García, V., Molina, F. P., Allen-Perkins, A., Godoy, O., & Bartomeus, I. (2026). Plant–pollinator interaction rewiring boosts year-to-year community persistence. Ecology Letters, 29 (1), e70293.

Dormann, C. F., Fründ, J., & Schaefer, H. M. (2017). Identifying causes of patterns in ecological networks: Opportunities and limitations. Annual Review of Ecology, Evolution, and Systematics, 48 (1), 559–584.

Dormann, C. F., Gruber, B., & Fründ, J. (2008). Introducing the bipartite package: Analysing ecological networks. Interaction, 1, 8–11.

Duchenne, F., Domínguez-García, V., Molina, F. P., & Bartomeus, I. (2026). Coevolution of phenological traits shapes plant–pollinator coexistence. Proceedings of the Royal Society B: Biological Sciences, 293 (2064).

Eklöf, A., Jacob, U., Kopp, J., Bosch, J., Castro-Urgal, R., Chacoff, N. P., Dalsgaard, B., de Sassi, C., Galetti, M., Guimarães, P. R., Lomáscolo, S. B., Martín González, A. M., Pizo, M. A., Rader, R., Rodrigo, A., Tylianakis, J. M., Vázquez, D. P., & Allesina, S. (2013). The dimensionality of ecological networks. Ecology Letters, 16 (5), 577–583.

Encinas-Viso, F., Revilla, T. A., & Etienne, R. S. (2012). Phenology drives mutualistic network structure and diversity. Ecology Letters, 15 (3), 198–208.

E-Vojtkó, A., de Bello, F., Durka, W., Kühn, I., & Götzenberger, L. (2020). The neglected importance of floral traits in trait-based plant community assembly. Journal of Vegetation Science, 31 (4), 529–539.

Földesi, R., Howlett, B. G., Grass, I., & Batáry, P. (2021). Larger pollinators deposit more pollen on stigmas across multiple plant species—A meta-analysis. Journal of Applied Ecology, 58 (4), 699–707.

Fort, H., Vázquez, D. P., & Lan, B. L. (2016). Abundance and generalisation in mutualistic networks: Solving the chicken-and-egg dilemma. Ecology Letters, 19 (1), 4–11.

Gómez-Martínez, C., González-Estévez, M. A., Cursach, J., & Lázaro, A. (2022). Pollinator richness, pollination networks, and diet adjustment along local and landscape gradients of resource diversity. Ecological Applications, 32 (6), e2634.

Greenwell, B., & Greenwell, M. B. (2020). fastshap (R package version 0.0.7). https://CRAN.R-project.org/package=fastshap

Guimarães, P. R., Jr. (2020). The structure of ecological networks across levels of organization. Annual Review of Ecology, Evolution, and Systematics, 51 (1), 433–460.

He, Y.-D., Lázaro, A., Bergamo, P. J., Liang, H., Yang, C.-F., & Ye, Z.-M. (2025). Disentangling the mechanisms behind indirect interactions between plants via shared pollinators: Effects of neutral and niche-based processes. Journal of Ecology, 113 (12), 3622–3636.

Jordano, P. (2016). Sampling networks of ecological interactions. Functional Ecology, 30 (12), 1883–1893.

Junker, R. R., Blüthgen, N., Brehm, T., Binkenstein, J., Paulus, J., Schaefer, H. M., & Stang, M. (2013). Specialization on traits as basis for the niche breadth of flower visitors and as structuring mechanism of ecological networks. Functional Ecology, 27 (2), 329–341.

Kaiser-Bunbury, C. N., Vázquez, D. P., Stang, M., & Ghazoul, J. (2014). Determinants of the microstructure of plant–pollinator networks. Ecology, 95 (12), 3314–3324.

Kelly, T. T., & Elle, E. (2021). Investigating bee dietary preferences along a gradient of floral resources: How does resource use align with resource availability? Insect Science, 28 (2), 555–565.

Kendall, L. K., Rader, R., Gagic, V., Cariveau, D. P., Albrecht, M., Baldock, K. C., Freitas, B. M., Hall, M., Holzschuh, A., Molina, F. P., & others. (2019). Pollinator size and its consequences: Robust estimates of body size in pollinating insects. Ecology and Evolution, 9 (4), 1702–1714.

Klumpers, S. G., Stang, M., & Klinkhamer, P. G. (2019). Foraging efficiency and size matching in a plant– pollinator community: The importance of sugar content and tongue length. Ecology Letters, 22 (3), 469–479.

Kudo, G., & Cooper, E. J. (2019). When spring ephemerals fail to meet pollinators: Mechanism of phenological mismatch and its impact on plant reproduction. Proceedings of the Royal Society B: Biological Sciences, 286 (1904).

Kuppler, J., Höfers, M. K., Wiesmann, L., & Junker, R. R. (2016). Time-invariant differences between plant individuals in interactions with arthropods correlate with intraspecific variation in plant phenology, morphology and floral scent. New Phytologist, 210 (4), 1357–1368.

Lanuza, J. B., Rader, R., Stavert, J., Kendall, L. K., Saunders, M. E., & Bartomeus, I. (2023). Covariation among reproductive traits in flowering plants shapes their interactions with pollinators. Functional Ecology, 37 (7), 2072–2084.

Leong, M., Ponisio, L. C., Kremen, C., Thorp, R. W., & Roderick, G. K. (2016). Temporal dynamics influenced by global change: Bee community phenology in urban, agricultural, and natural landscapes. Global Change Biology, 22 (3), 1046–1053.

Morán-López, T., Benadi, G., Lara-Romero, C., Chacoff, N., Vitali, A., Pescador, D., Lomáscolo, S. B., Morente-López, J., Vázquez, D. P., & Morales, J. M. (2022). Flexible diets enable pollinators to cope with changes in plant community composition. Journal of Ecology, 110 (8), 1913–1927.

Nordt, B., Hensen, I., Bucher, S. F., Freiberg, M., Primack, R. B., Stevens, A.-D., Bonn, A., Wirth, C., Jakubka, D., Plos, C., & others. (2021). The PhenObs initiative: A standardised protocol for monitoring phenological responses to climate change using herbaceous plant species in botanical gardens. Functional Ecology, 35 (4), 821–834.

Oksanen, J., Simpson, G., Blanchet, F., Kindt, R., Legendre, P., Minchin, P., O’Hara, R., Solymos, P., Stevens, M., Szoecs, E., Wagner, H., Bedward, M., Bolker, B., Borcard, D., Carvalho, G., De Caceres, M., Durand, S., Evangelista, H., Hannigan, G., Hill, M., Lahti, L., Martino, C., Ouellette, M., Ribeiro Cunha, E., Smith, T., Stier, A., Ter Braak, C., & Weedon, J. (2026). vegan: Community Ecology Package (R package version 2. 8–0). https://vegandevs.github.io/vegan/

Olesen, J. M., Bascompte, J., Dupont, Y. L., Elberling, H., Rasmussen, C., & Jordano, P. (2011). Missing and forbidden links in mutualistic networks. Proceedings of the Royal Society B: Biological Sciences, 278 (1706), 725–732.

Olito, C., & Fox, J. W. (2015). Species traits and abundances predict metrics of plant–pollinator network structure, but not pairwise interactions. Oikos, 124 (4), 428–436.

Payrató-Borràs, C., Gracia-Lázaro, C., Hernández, L., & Moreno, Y. (2024). Beyond the aggregated paradigm: Phenology and structure in mutualistic networks. Journal of Physics: Complexity, 5 (2), 025013.

Peralta, G., CaraDonna, P. J., Rakosy, D., Fründ, J., Tudanca, M. P. P., Dormann, C. F., Burkle, L. A., Kaiser-Bunbury, C. N., Knight, T. M., Resasco, J., & others. (2024). Predicting plant–pollinator interactions: Concepts, methods, and challenges. Trends in Ecology & Evolution, 39 (5), 494–505.

Peralta, G., Perry, G. L., Vázquez, D. P., Dehling, D. M., & Tylianakis, J. M. (2020). Strength of niche processes for species interactions is lower for generalists and exotic species. Journal of Animal Ecology, 89 (9), 2145–2155.

Peralta, G., Vázquez, D. P., Chacoff, N. P., Lomáscolo, S. B., Perry, G. L. W., & Tylianakis, J. M. (2020). Trait matching and phenological overlap increase the spatio-temporal stability and functionality of plant– pollinator interactions. Ecology Letters, 23 (7), 1107–1116.

Pichler, M., Boreux, V., Klein, A.-M., Schleuning, M., & Hartig, F. (2020). Machine learning algorithms to infer trait-matching and predict species interactions in ecological networks. Methods in Ecology and Evolution, 11 (2), 281–293.

Poisot, T., Stouffer, D. B., & Gravel, D. (2015). Beyond species: Why ecological interaction networks vary through space and time. Oikos, 124 (3), 243–251.

R Core Team. (2025). R: A language and environment for statistical computing. R Foundation for Statistical Computing. https://www.R-project.org/

Resasco, J., Chacoff, N. P., & Vázquez, D. P. (2021). Plant–pollinator interactions between generalists persist over time and space. Ecology, 102 (6), e03359.

Rocha-Ortega, M., Rodriguez, P., & Córdoba-Aguilar, A. (2021). Geographical, temporal and taxonomic biases in insect GBIF data on biodiversity and extinction. Ecological Entomology, 46 (4), 718–728.

Sánchez-Martín, R., Barreto, E., Maxwell, M. F., Duchenne, F., Beck, H., Bobato, R., Brenes, E., Bôlla, D., Büttner, N., Caron, A. P., & others. (2025). Mechanisms influencing network topology in plant– hummingbird pollination networks. Proceedings of the Royal Society B: Biological Sciences, 292 (2059).

Sazatornil, F. D., More, M., Benitez-Vieyra, S., Cocucci, A. A., Kitching, I. J., Schlumpberger, B. O., Oliveira, P. E., Sazima, M., & Amorim, F. W. (2016). Beyond neutral and forbidden links: Morphological matches and the assembly of mutualistic hawkmoth–plant networks. Journal of Animal Ecology, 85 (6), 1586–1594.

Schemske, D. W., Mittelbach, G. G., Cornell, H. V., Sobel, J. M., & Roy, K. (2009). Is there a latitudinal gradient in the importance of biotic interactions? Annual Review of Ecology, Evolution, and Systematics, 40 (1), 245–269.

Schwarz, B., Vázquez, D. P., CaraDonna, P. J., Knight, T. M., Benadi, G., Dormann, C. F., Gauzens, B., Motivans, E., Resasco, J., Blüthgen, N., & others. (2020). Temporal scale-dependence of plant–pollinator networks. Oikos, 129 (9), 1289–1302.

Simmons, B. I., Balmford, A., Bladon, A. J., Christie, A. P., De Palma, A., Dicks, L. V., Gallego-Zamorano, J., Johnston, A., Martin, P. A., Purvis, A., & others. (2019). Worldwide insect declines: An important message, but interpret with caution. Ecology and Evolution, 9 (7), 3678–3680.

Stang, M., Klinkhamer, P. G., & Van der Meijden, E. (2007). Asymmetric specialization and extinction risk in plant–flower visitor webs: A matter of morphology or abundance? Oecologia, 151 (3), 442–453.

Strydom, T., Catchen, M. D., Banville, F., Caron, D., Dansereau, G., Desjardins-Proulx, P., … & Poisot, T. (2021). A roadmap towards predicting species interaction networks (across space and time). Philosophical Transactions of the Royal Society B: Biological Sciences, 376 (1837), 20210063.

Sweet, F. S., Apfelbeck, B., Hanusch, M., Garland Monteagudo, C., & Weisser, W. W. (2022). Data from public and governmental databases show that a large proportion of the regional animal species pool occur in cities in Germany. Journal of Urban Ecology, 8 (1), juac002.

Tylianakis, J. M., & Morris, R. J. (2017). Ecological networks across environmental gradients. Annual Review of Ecology, Evolution, and Systematics, 48 (1), 25–48.

Valdovinos, F. S. (2019). Mutualistic networks: Moving closer to a predictive theory. Ecology Letters, 22 (9), 1517–1534.

Vázquez, D. P., Blüthgen, N., Cagnolo, L., & Chacoff, N. P. (2009). Uniting pattern and process in plant– animal mutualistic networks: A review. Annals of Botany, 103 (9), 1445–1457.

Vázquez, D. P., Lomáscolo, S. B., Maldonado, M. B., Chacoff, N. P., Dorado, J., Stevani, E. L., & Vitale, N. L. (2012). The strength of plant–pollinator interactions. Ecology, 93 (4), 719–725.

Vázquez, D. P., Vitale, N., Dorado, J., Amico, G., & Stevani, E. L. (2023). Phenological mismatches and the demography of solitary bees. Proceedings of the Royal Society B: Biological Sciences, 290 (1990).

Vizentin-Bugoni, J., Maruyama, P. K., & Sazima, M. (2014). Processes entangling interactions in communities: Forbidden links are more important than abundance in a hummingbird–plant network. Proceedings of the Royal Society B: Biological Sciences, 281 (1780), 20132397.

Williams, R. J., Anandanadesan, A., & Purves, D. (2010). The probabilistic niche model reveals the niche structure and role of body size in a complex food web. PLoS ONE, 5 (8), e12092.

Wood, S. (2015). mgcv: Mixed GAM computation vehicle with automatic smoothness estimation (R package).

Wright, M. N., Wager, S., Probst, P., & Wright, M. M. N. (2019). ranger: A fast implementation of random forests (R package, Version 0.11.2).

Zhao, Y.-H., Lázaro, A., Li, H.-D., Tao, Z.-B., Liang, H., Zhou, W., Ren, Z.-X., Xu, K., Li, D.-Z., & Wang, H. (2022). Morphological trait-matching in plant–Hymenoptera and plant–Diptera mutualisms across an elevational gradient. Journal of Animal Ecology, 91 (1), 196–209.

