## Supplementary Material for "Seasonal Variation Mediates the Importance of Species Attributes in Plant-Pollinator Interactions"

**Table S1.** List of focal plant species included in the study, along with their family and order.

| Species | Family | Order |
|---|---|---|
| *Cyclamen coum* | Primulaceae | Ericales |
| *Helleborus foetidus* | Ranunculaceae | Ranunculales |
| *Tussilago farfara* | Asteraceae | Asterales |
| *Vinca minor* | Apocynaceae | Gentianales |
| *Scopolia carniolica* | Solanaceae | Solanales |
| *Viola odorata* | Violaceae | Malpighiales |
| *Puschkinia scilloides* | Asparagaceae | Asparagales |
| *Lathyrus vernus* | Fabaceae | Fabales |
| *Hepatica nobilis* | Ranunculaceae | Ranunculales |
| *Pulsatilla vulgaris* | Ranunculaceae | Ranunculales |
| *Primula denticulata* | Primulaceae | Ericales |
| *Narcissus pseudonarcissus* | Amaryllidaceae | Asparagales |
| *Primula veris* | Primulaceae | Ericales |
| *Iberis sempervirens* | Brassicaceae | Brassicales |
| *Anemone nemorosa* | Ranunculaceae | Ranunculales |
| *Trillium sessile* | Melanthiaceae | Liliales |
| *Caltha palustris* | Ranunculaceae | Ranunculales |
| *Bergenia purpurascens* | Saxifragaceae | Saxifragales |
| *Geum rivale* | Rosaceae | Rosales |
| *Viscaria vulgaris* | Caryophyllaceae | Caryophyllales |
| *Tulipa sylvestris* | Liliaceae | Liliales |
| *Fragaria vesca* | Rosaceae | Rosales |
| *Oxalis acetosella* | Oxalidaceae | Oxalidales |
| *Brunnera macrophylla* | Boraginaceae | Boraginales |
| *Lamium album* | Lamiaceae | Lamiales |
| *Paeonia officinalis* | Paeoniaceae | Saxifragales |
| *Chelidonium majus* | Papaveraceae | Ranunculales |
| *Polygonum bistorta* | Polygonaceae | Caryophyllales |
| *Anemone sylvestris* | Ranunculaceae | Ranunculales |
| *Asarum caudatum* | Aristolochiaceae | Piperales |
| *Dryas octopetala* | Rosaceae | Rosales |
| *Aquilegia vulgaris* | Ranunculaceae | Ranunculales |
| *Allium ursinum* | Amaryllidaceae | Asparagales |
| *Paeonia delavayi* | Paeoniaceae | Saxifragales |
| *Menyanthes trifoliata* | Menyanthaceae | Asterales |
| *Convallaria majalis* | Asparagaceae | Asparagales |
| *Helianthemum nummularium* | Cistaceae | Malvales |
| *Geranium sanguineum* | Geraniaceae | Geraniales |
| *Asphodelus albus* | Asphodelaceae | Asparagales |
| *Linum perenne* | Linaceae | Malpighiales |
| *Vincetoxicum hirundinaria* | Apocynaceae | Gentianales |
| *Psephellus dealbatus* | Asteraceae | Asterales |
| *Aconitum lycoctonum* | Ranunculaceae | Ranunculales |

| | | |
|---|---|---|
| *Dictamnus albus* | Rutaceae | Sapindales |
| *Silene nutans* | Caryophyllaceae | Caryophyllales |
| *Salvia pratensis* | Lamiaceae | Lamiales |
| *Salvia officinalis* | Lamiaceae | Lamiales |
| *Campanula rotundifolia* | Campanulaceae | Asterales |
| *Lotus maritimus* | Fabaceae | Fabales |
| *Solanum dulcamara* | Solanaceae | Solanales |
| *Centranthus ruber* | Caprifoliaceae | Dipsacales |
| *Gratiola officinalis* | Plantaginaceae | Lamiales |
| *Lotus corniculatus* | Fabaceae | Fabales |
| *Hemerocallis dumortieri* | Asphodelaceae | Asparagales |
| *Silene vulgaris* | Caryophyllaceae | Caryophyllales |
| *Hypericum olympicum* | Hypericaceae | Malpighiales |
| *Leucanthemum vulgare* | Asteraceae | Asterales |
| *Achillea millefolium* | Asteraceae | Asterales |
| *Penstemon bradburyi* | Plantaginaceae | Lamiales |
| *Clematis recta* | Ranunculaceae | Ranunculales |
| *Lavandula angustifolia* | Lamiaceae | Lamiales |
| *Hypericum perforatum* | Hypericaceae | Malpighiales |
| *Silene coronaria* | Caryophyllaceae | Caryophyllales |
| *Anthericum ramosum* | Asparagaceae | Asparagales |
| *Coronilla varia* | Fabaceae | Fabales |
| *Galega officinalis* | Fabaceae | Fabales |
| *Genista tinctoria* | Fabaceae | Fabales |
| *Betonica officinalis* | Lamiaceae | Lamiales |
| *Platycodon grandiflorus* | Campanulaceae | Asterales |
| *Hemerocallis fulva* | Asphodelaceae | Asparagales |
| *Clematis integrifolia* | Ranunculaceae | Ranunculales |
| *Dianthus carthusianorum* | Caryophyllaceae | Caryophyllales |
| *Saponaria officinalis* | Caryophyllaceae | Caryophyllales |
| *Fuchsia magellanica* | Onagraceae | Myrtales |
| *Eriocapitella hupehensis* | Ranunculaceae | Ranunculales |
| *Tanacetum vulgare* | Asteraceae | Asterales |
| *Origanum vulgare* | Lamiaceae | Lamiales |
| *Scabiosa ochroleuca* | Caprifoliaceae | Dipsacales |

**Table S2.** First and last sampling dates and total number of weekly surveys conducted in each botanical garden during the 2023 flowering season.

| Botanical garden | Start | End | Total surveys |
|---|---|---|---|
| Halle | 29 Mar | 23 Aug | 21 |
| Jena | 31 Mar | 18 Aug | 19 |
| Leipzig | 23 Mar | 23 Aug | 22 |

**Table S3.** Focal plant species included in the study, with sampling time (minutes) per botanical garden (Leipzig, Halle, and Jena). The table also reports the total sampling time per species across all gardens. Species sampled in all three gardens are highlighted in bold. Total sampling time was 49 h (Leipzig), 52 h (Halle), and 40 h (Jena), summing to 141 h overall.

| Species | Leipzig (min) | Halle (min) | Jena (min) | Total time (min) |
|---|---|---|---|---|
| *Achillea millefolium* | 0 | 90 | 93 | 183 |
| *Aconitum lycoctonum* | 0 | 75 | 0 | 75 |
| *Allium ursinum* | 30 | 0 | 45 | 75 |
| ***Anemone nemorosa*** | 45 | 45 | 45 | 135 |
| *Anemone sylvestris* | 30 | 90 | 0 | 120 |
| *Anthericum ramosum* | 75 | 0 | 0 | 75 |
| *Aquilegia vulgaris* | 165 | 0 | 30 | 195 |
| *Asarum caudatum* | 16 | 0 | 0 | 16 |
| *Asphodelus albus* | 0 | 0 | 30 | 30 |
| *Bergenia purpurascens* | 75 | 0 | 45 | 120 |
| *Betonica officinalis* | 0 | 0 | 45 | 45 |
| *Brunnera macrophylla* | 0 | 0 | 105 | 105 |
| *Caltha palustris* | 120 | 0 | 0 | 120 |
| *Campanula rotundifolia* | 0 | 0 | 75 | 75 |
| *Centranthus ruber* | 150 | 150 | 0 | 300 |
| *Chelidonium majus* | 0 | 90 | 75 | 165 |
| *Clematis integrifolia* | 0 | 105 | 0 | 105 |
| *Clematis recta* | 0 | 105 | 15 | 120 |
| *Convallaria majalis* | 45 | 0 | 0 | 45 |
| *Coronilla varia* | 0 | 0 | 63 | 63 |
| *Cyclamen coum* | 30 | 0 | 0 | 30 |
| *Dianthus carthusianorum* | 0 | 91 | 0 | 91 |
| ***Dictamnus albus*** | 45 | 45 | 30 | 120 |
| *Dryas octopetala* | 60 | 0 | 0 | 60 |
| *Eriocapitella hupehensis* | 90 | 0 | 75 | 165 |
| *Fragaria vesca* | 75 | 150 | 0 | 225 |
| *Fuchsia magellanica* | 0 | 105 | 18 | 123 |
| *Galega officinalis* | 0 | 0 | 78 | 78 |
| *Genista tinctoria* | 0 | 0 | 33 | 33 |
| *Geranium sanguineum* | 75 | 0 | 90 | 165 |
| *Geum rivale* | 0 | 90 | 60 | 150 |
| *Gratiola officinalis* | 0 | 30 | 0 | 30 |
| *Helianthemum nummularium* | 75 | 75 | 0 | 150 |
| *Helleborus foetidus* | 120 | 0 | 15 | 135 |
| *Hemerocallis dumortieri* | 45 | 0 | 0 | 45 |
| *Hemerocallis fulva* | 0 | 30 | 0 | 30 |
| *Hepatica nobilis* | 0 | 0 | 80 | 80 |
| ***Hypericum olympicum*** | 105 | 45 | 30 | 180 |
| *Hypericum perforatum* | 0 | 45 | 33 | 78 |
| ***Iberis sempervirens*** | 120 | 105 | 105 | 330 |
| ***Lamium album*** | 15 | 105 | 105 | 225 |

| | | | | |
|---|---|---|---|---|
| ***Lathyrus vernus*** | 60 | 180 | 45 | 285 |
| ***Lavandula angustifolia*** | 60 | 75 | 93 | 228 |
| *Leucanthemum vulgare* | 0 | 15 | 33 | 48 |
| *Linum perenne* | 0 | 0 | 78 | 78 |
| *Lotus corniculatus* | 0 | 135 | 0 | 135 |
| *Lotus maritimus* | 0 | 0 | 30 | 30 |
| *Menyanthes trifoliata* | 30 | 0 | 0 | 30 |
| *Narcissus pseudonarcissus* | 0 | 45 | 0 | 45 |
| *Origanum vulgare* | 0 | 45 | 75 | 120 |
| *Oxalis acetosella* | 0 | 15 | 0 | 15 |
| *Paeonia delavayi* | 0 | 0 | 45 | 45 |
| *Paeonia officinalis* | 30 | 0 | 30 | 60 |
| *Penstemon bradburyi* | 60 | 0 | 0 | 60 |
| *Platycodon grandiflorus* | 30 | 75 | 0 | 105 |
| *Polygonum bistorta* | 0 | 75 | 0 | 75 |
| ***Primula denticulata*** | 60 | 30 | 90 | 180 |
| ***Primula veris*** | 60 | 75 | 45 | 180 |
| *Psephellus dealbatus* | 90 | 0 | 0 | 90 |
| ***Pulsatilla vulgaris*** | 105 | 15 | 60 | 180 |
| *Puschkinia scilloides* | 0 | 45 | 0 | 45 |
| *Salvia officinalis* | 30 | 60 | 0 | 90 |
| *Salvia pratensis* | 0 | 0 | 45 | 45 |
| *Saponaria officinalis* | 0 | 105 | 0 | 105 |
| *Scabiosa ochroleuca* | 45 | 0 | 0 | 45 |
| *Scopolia carniolica* | 91 | 105 | 0 | 196 |
| *Silene coronaria* | 30 | 0 | 0 | 30 |
| *Silene nutans* | 0 | 0 | 30 | 30 |
| *Silene vulgaris* | 75 | 0 | 0 | 75 |
| *Solanum dulcamara* | 0 | 75 | 0 | 75 |
| *Tanacetum vulgare* | 45 | 60 | 0 | 105 |
| *Trillium sessile* | 76 | 0 | 60 | 136 |
| *Tulipa sylvestris* | 0 | 15 | 0 | 15 |
| *Tussilago farfara* | 103 | 0 | 0 | 103 |
| *Vinca minor* | 150 | 0 | 75 | 225 |
| ***Vincetoxicum hirundinaria*** | 75 | 90 | 90 | 255 |
| *Viola odorata* | 0 | 60 | 0 | 60 |
| ***Viscaria vulgaris*** | 30 | 75 | 60 | 165 |

**Table S4** List of non-focal plant species included in the study, along with their family and order.

| Species | Family | Order |
|---|---|---|
| *Achillea filipendulina* | Asteraceae | Asterales |
| *Aconitum lycoctonum* | Ranunculaceae | Ranunculales |
| *Aegonychon purpurocaeruleum* | Boraginaceae | Boraginales |
| *Allium cernuum* | Amaryllidaceae | Asparagales |
| *Allium tuberosum* | Amaryllidaceae | Asparagales |
| *Allium turkestanicum* | Amaryllidaceae | Asparagales |
| *Alyssum saxatile* | Brassicaceae | Brassicales |
| *Anthriscus sylvestris* | Apiaceae | Apiales |
| *Arabis alpina* | Brassicaceae | Brassicales |
| *Asclepias syriaca* | Apocynaceae | Gentianales |
| *Asparagus oligoclonos* | Asparagaceae | Asparagales |
| *Astragalus alopecurus* | Fabaceae | Fabales |
| *Aubrieta deltoidea* | Brassicaceae | Brassicales |
| *Aubrieta olympica* | Brassicaceae | Brassicales |
| *Aubrieta parviflora* | Brassicaceae | Brassicales |
| *Aubrieta pinardii* | Brassicaceae | Brassicales |
| *Ballota nigra* | Lamiaceae | Lamiales |
| *Baptisia australis* | Fabaceae | Fabales |
| *Barbarea vulgaris* | Brassicaceae | Brassicales |
| *Bellis perennis* | Asteraceae | Asterales |
| *Bupleurum aureum* | Apiaceae | Apiales |
| *Campanula poscharskyana* | Campanulaceae | Asterales |
| *Centaurea cyanus* | Asteraceae | Asterales |
| *Cephalaria uralensis* | Caprifoliaceae | Dipsacales |
| *Cerinthe minor* | Boraginaceae | Boraginales |
| *Chrysojasminum fruticans* | Oleaceae | Lamiales |
| *Cistus salviifolius* | Cistaceae | Malvales |
| *Clinopodium nepeta* | Lamiaceae | Lamiales |
| *Corydalis cava* | Papaveraceae | Ranunculales |
| *Cosmos sulphureus* | Asteraceae | Asterales |
| *Cotoneaster adpressus* | Rosaceae | Rosales |
| *Crambe maritima* | Brassicaceae | Brassicales |
| *Cylindropuntia imbricata* | Cactaceae | Caryophyllales |
| *Daucus carota* | Apiaceae | Apiales |
| *Delosperma cooperi* | Aizoaceae | Caryophyllales |
| *Delphinium elatum* | Ranunculaceae | Ranunculales |
| *Digitalis purpurea* | Plantaginaceae | Lamiales |
| *Drymocallis rupestris* | Rosaceae | Rosales |
| *Erodium cicutarium* | Geraniaceae | Geraniales |
| *Eryngium caeruleum* | Apiaceae | Apiales |
| *Eryngium maritimum* | Apiaceae | Apiales |
| *Eryngium planum* | Apiaceae | Apiales |
| *Erysimum pulchellum* | Brassicaceae | Brassicales |
| *Eschscholzia californica* | Papaveraceae | Ranunculales |
| *Ferula assa-foetida* | Apiaceae | Apiales |
| *Foeniculum vulgare* | Apiaceae | Apiales |
| *Genista hispanica* | Fabaceae | Fabales |
| *Genista radiata* | Fabaceae | Fabales |
| *Geum ternatum* | Rosaceae | Rosales |
| *Globularia trichosantha* | Plantaginaceae | Lamiales |

| | | |
|---|---|---|
| *Gypsophila paniculata* | Caryophyllaceae | Caryophyllales |
| *Helianthemum apenninum* | Cistaceae | Malvales |
| *Heuchera grossulariifolia* | Saxifragaceae | Saxifragales |
| *Hippocrepis emerus* | Fabaceae | Fabales |
| *Hippomarathrum vulgare* | Apiaceae | Apiales |
| *Hypochaeris radicata* | Asteraceae | Asterales |
| *Impatiens glandulifera* | Balsaminaceae | Ericales |
| *Inula helenium* | Asteraceae | Asterales |
| *Ipheion uniflorum* | Amaryllidaceae | Asparagales |
| *Knautia dipsacifolia* | Caprifoliaceae | Dipsacales |
| *Lamium orvala* | Lamiaceae | Lamiales |
| *Lamium purpureum* | Lamiaceae | Lamiales |
| *Liatris pycnostachya* | Asteraceae | Asterales |
| *Limnanthes douglasii* | Limnanthaceae | Brassicales |
| *Limonium gmelini* | Plumbaginaceae | Caryophyllales |
| *Limonium sinuatum* | Plumbaginaceae | Caryophyllales |
| *Lythrum salicaria* | Lythraceae | Myrtales |
| *Magnolia stellata* | Magnoliaceae | Magnoliales |
| *Malcolmia maritima* | Brassicaceae | Brassicales |
| *Malva thuringiaca* | Malvaceae | Malvales |
| *Marrubium peregrinum* | Lamiaceae | Lamiales |
| *Melilotus albus* | Fabaceae | Fabales |
| *Monarda citriodora* | Lamiaceae | Lamiales |
| *Moricandia arvensis* | Brassicaceae | Brassicales |
| *Muscari armeniacum* | Asparagaceae | Asparagales |
| *Muscari botryoides* | Asparagaceae | Asparagales |
| *Muscari comosum* | Asparagaceae | Asparagales |
| *Nepeta clarkei* | Lamiaceae | Lamiales |
| *Oenothera odorata* | Onagraceae | Myrtales |
| *Ornithogalum nutans* | Asparagaceae | Asparagales |
| *Paeonia daurica* | Paeoniaceae | Saxifragales |
| *Penstemon hartwegii* | Plantaginaceae | Lamiales |
| *Pentaglottis sempervirens* | Boraginaceae | Boraginales |
| *Phacelia congesta* | Hydrophyllaceae | Boraginales |
| *Phedimus kamtschaticus* | Crassulaceae | Saxifragales |
| *Phlomoides tuberosa* | Lamiaceae | Lamiales |
| *Pieris japonica* | Ericaceae | Ericales |
| *Potentilla delphinensis* | Rosaceae | Rosales |
| *Potentilla montana* | Rosaceae | Rosales |
| *Potentilla sterilis* | Rosaceae | Rosales |
| *Primula vulgaris* | Primulaceae | Ericales |
| *Prunella grandiflora* | Lamiaceae | Lamiales |
| *Prunus tenella* | Rosaceae | Rosales |
| *Pulmonaria mollis* | Boraginaceae | Boraginales |
| *Ranunculus bulbosus* | Ranunculaceae | Ranunculales |
| *Ranunculus caucasicus* | Ranunculaceae | Ranunculales |
| *Ranunculus ficaria* | Ranunculaceae | Ranunculales |
| *Ranunculus lanuginosus* | Ranunculaceae | Ranunculales |
| *Reseda alba* | Resedaceae | Brassicales |
| *Reseda lutea* | Resedaceae | Brassicales |
| *Reseda odorata* | Resedaceae | Brassicales |
| *Rhododendron smirnowii* | Ericaceae | Ericales |
| *Rosa foetida* | Rosaceae | Rosales |
| *Rudbeckia laciniata* | Asteraceae | Asterales |
| *Salvia sclarea* | Lamiaceae | Lamiales |
| *Satureja montana* | Lamiaceae | Lamiales |

| | | |
|---|---|---|
| *Scutellaria baicalensis* | Lamiaceae | Lamiales |
| *Seseli gummiferum* | Apiaceae | Apiales |
| *Silene chungtienensis* | Caryophyllaceae | Caryophyllales |
| *Sisymbrium officinale* | Brassicaceae | Brassicales |
| *Stachys byzantina* | Lamiaceae | Lamiales |
| *Succisella inflexa* | Caprifoliaceae | Dipsacales |
| *Symphytum officinale* | Boraginaceae | Boraginales |
| *Syringa tomentella* | Oleaceae | Lamiales |
| *Tagetes erecta* | Asteraceae | Asterales |
| *Taraxacum officinale* | Asteraceae | Asterales |
| *Teucrium scorodonia* | Lamiaceae | Lamiales |
| *Thalictrum flavum* | Ranunculaceae | Ranunculales |
| *Thymus vulgaris* | Lamiaceae | Lamiales |
| *Trifolium rubens* | Fabaceae | Fabales |
| *Valeriana phu* | Caprifoliaceae | Dipsacales |
| *Verbena rigida* | Verbenaceae | Lamiales |
| *Wisteria sinensis* | Fabaceae | Fabales |
| *Zinnia elegans* | Asteraceae | Asterales |
| *Acantholimon ulicinum* | Plumbaginaceae | Caryophyllales |
| *Agapanthus africanus* | Amaryllidaceae | Asparagales |
| *Alkanna orientalis* | Boraginaceae | Boraginales |
| *Anemone ranunculoides* | Ranunculaceae | Ranunculales |
| *Anthyllis vulneraria* | Fabaceae | Fabales |
| *Bergenia ciliata* | Saxifragaceae | Saxifragales |
| *Betonica alopecuros* | Lamiaceae | Lamiales |
| *Borago officinalis* | Boraginaceae | Boraginales |
| *Brassica oleracea* | Brassicaceae | Brassicales |
| *Buddleja davidii* | Scrophulariaceae | Lamiales |
| *Campanula alliariifolia* | Campanulaceae | Asterales |
| *Campanula glomerata* | Campanulaceae | Asterales |
| *Campanula lactiflora* | Campanulaceae | Asterales |
| *Campanula persicifolia* | Campanulaceae | Asterales |
| *Campanula pyramidalis* | Campanulaceae | Asterales |
| *Campanula rapunculoides* | Campanulaceae | Asterales |
| *Cardamine pratensis* | Brassicaceae | Brassicales |
| *Centaurea scabiosa* | Asteraceae | Asterales |
| *Centaurea stoebe* | Asteraceae | Asterales |
| *Centaurea subtilis* | Asteraceae | Asterales |
| *Cerastium arvense* | Caryophyllaceae | Caryophyllales |
| *Chamaecytisus austriacus* | Fabaceae | Fabales |
| *Chamaecytisus hirsutus* | Fabaceae | Fabales |
| *Cichorium intybus* | Asteraceae | Asterales |
| *Coronilla orientalis* | Fabaceae | Fabales |
| *Cytisophyllum sessilifolium* | Fabaceae | Fabales |
| *Dasiphora fruticosa* | Rosaceae | Rosales |
| *Desmodium canadense* | Fabaceae | Fabales |
| *Dianthus petraeus* | Caryophyllaceae | Caryophyllales |
| *Digitalis grandiflora* | Plantaginaceae | Lamiales |
| *Digitalis obscura* | Plantaginaceae | Lamiales |
| *Dracocephalum rupestre* | Lamiaceae | Lamiales |
| *Erigeron annuus* | Asteraceae | Asterales |
| *Euphorbia nicaeensis* | Euphorbiaceae | Malpighiales |
| *Eutrochium maculatum* | Asteraceae | Asterales |
| *Fagopyrum esculentum* | Polygonaceae | Caryophyllales |
| *Genista germanica* | Fabaceae | Fabales |
| *Geranium molle* | Geraniaceae | Geraniales |

| | | |
|---|---|---|
| *Globularia cordifolia* | Plantaginaceae | Lamiales |
| *Helichrysum arenarium* | Asteraceae | Asterales |
| *Heracleum sphondylium* | Apiaceae | Apiales |
| *Heuchera villosa* | Saxifragaceae | Saxifragales |
| *Hypericum fragile* | Hypericaceae | Malpighiales |
| *Hypericum hookerianum* | Hypericaceae | Malpighiales |
| *Inula salicina* | Asteraceae | Asterales |
| *Iris germanica* | Iridaceae | Asparagales |
| *Iris pseudacorus* | Iridaceae | Asparagales |
| *Jacobaea erucifolia* | Asteraceae | Asterales |
| *Knautia arvensis* | Caprifoliaceae | Dipsacales |
| *Lactuca macrophylla* | Asteraceae | Asterales |
| *Lactuca plumieri* | Asteraceae | Asterales |
| *Lamium galeobdolon* | Lamiaceae | Lamiales |
| *Lamprocapnos spectabilis* | Papaveraceae | Ranunculales |
| *Lathyrus aureus* | Fabaceae | Fabales |
| *Lathyrus niger* | Fabaceae | Fabales |
| *Lathyrus pratensis* | Fabaceae | Fabales |
| *Lilium martagon* | Liliaceae | Liliales |
| *Lindelofia anchusoides* | Boraginaceae | Boraginales |
| *Lomelosia albocincta* | Caprifoliaceae | Dipsacales |
| *Lonicera floribunda* | Caprifoliaceae | Dipsacales |
| *Lysimachia vulgaris* | Primulaceae | Ericales |
| *Medicago sativa* | Fabaceae | Fabales |
| *Omphalodes verna* | Boraginaceae | Boraginales |
| *Paeonia mascula* | Paeoniaceae | Saxifragales |
| *Papaver nudicaule* | Papaveraceae | Ranunculales |
| *Papaver somniferum* | Papaveraceae | Ranunculales |
| *Pentanema helenioides* | Asteraceae | Asterales |
| *Phacelia tanacetifolia* | Hydrophyllaceae | Boraginales |
| *Phedimus aizoon* | Crassulaceae | Saxifragales |
| *Philadelphus pekinensis* | Hydrangeaceae | Cornales |
| *Phlox paniculata* | Polemoniaceae | Ericales |
| *Pilosella officinarum* | Asteraceae | Asterales |
| *Rhododendron saluenense* | Ericaceae | Ericales |
| *Rosa spinosissima* | Rosaceae | Rosales |
| *Rudbeckia fulgida* | Asteraceae | Asterales |
| *Salvia nutans* | Lamiaceae | Lamiales |
| *Satureja parnassica* | Lamiaceae | Lamiales |
| *Scabiosa canescens* | Caprifoliaceae | Dipsacales |
| *Scrophularia versicolor* | Scrophulariaceae | Lamiales |
| *Scutellaria altissima* | Lamiaceae | Lamiales |
| *Scutellaria cordifolia* | Lamiaceae | Lamiales |
| *Seseli montanum* | Apiaceae | Apiales |
| *Solidago flexicaulis* | Asteraceae | Asterales |
| *Spartium junceum* | Fabaceae | Fabales |
| *Spiraea chamaedryfolia* | Rosaceae | Rosales |
| *Stellaria palustris* | Caryophyllaceae | Caryophyllales |
| *Symphytum bulbosum* | Boraginaceae | Boraginales |
| *Teucrium canadense* | Lamiaceae | Lamiales |
| *Teucrium chamaedrys* | Lamiaceae | Lamiales |
| *Tilia henryana* | Malvaceae | Malvales |
| *Trifolium pratense* | Fabaceae | Fabales |
| *Tripleurospermum maritimum* | Asteraceae | Asterales |
| *Trommsdorffia maculata* | Asteraceae | Asterales |
| *Verbascum phoeniceum* | Scrophulariaceae | Lamiales |

| | | |
|---|---|---|
| *Vicia sepium* | Fabaceae | Fabales |
| *Aethionema grandiflorum* | Brassicaceae | Brassicales |
| *Ajuga reptans* | Lamiaceae | Lamiales |
| *Alisma plantago-aquatica* | Alismataceae | Alismatales |
| *Allium senescens* | Amaryllidaceae | Asparagales |
| *Anaphalis margaritacea* | Asteraceae | Asterales |
| *Anchusa officinalis* | Boraginaceae | Boraginales |
| *Aster mongolicus* | Asteraceae | Asterales |
| *Aurinia saxatilis* | Brassicaceae | Brassicales |
| *Carlina vulgaris* | Asteraceae | Asterales |
| *Centaurea nigra* | Asteraceae | Asterales |
| *Centaurea sadleriana* | Asteraceae | Asterales |
| *Cephalaria gigantea* | Caprifoliaceae | Dipsacales |
| *Cephalaria procera* | Caprifoliaceae | Dipsacales |
| *Chaenorhinum origanifolium* | Plantaginaceae | Lamiales |
| *Cirsium vulgare* | Asteraceae | Asterales |
| *Clinopodium album* | Lamiaceae | Lamiales |
| *Convolvulus tricolor* | Convolvulaceae | Solanales |
| *Coristospermum lucidum* | Apiaceae | Apiales |
| *Cuphea viscosissima* | Lythraceae | Myrtales |
| *Dasiphora davurica* | Rosaceae | Rosales |
| *Echinacea angustifolia* | Asteraceae | Asterales |
| *Echinops pungens* | Asteraceae | Asterales |
| *Echium vulgare* | Boraginaceae | Boraginales |
| *Erica carnea* | Ericaceae | Ericales |
| *Eriocapitella tomentosa* | Ranunculaceae | Ranunculales |
| *Erodium manescavi* | Geraniaceae | Geraniales |
| *Euphorbia characias* | Euphorbiaceae | Malpighiales |
| *Euphorbia seguieriana* | Euphorbiaceae | Malpighiales |
| *Ferulago aucheri* | Apiaceae | Apiales |
| *Forsythia intermedia* | Oleaceae | Lamiales |
| *Geranium macrorrhizum* | Geraniaceae | Geraniales |
| *Helichrysum italicum* | Asteraceae | Asterales |
| *Hieracium murorum* | Asteraceae | Asterales |
| *Impatiens balsamina* | Balsaminaceae | Ericales |
| *Inula oculus-christi* | Asteraceae | Asterales |
| *Klasea radiata* | Asteraceae | Asterales |
| *Knautia macedonica* | Caprifoliaceae | Dipsacales |
| *Lamium flexuosum* | Lamiaceae | Lamiales |
| *Lamium hybridum* | Lamiaceae | Lamiales |
| *Lathyrus roseus* | Fabaceae | Fabales |
| *Lomelosia cretica* | Caprifoliaceae | Dipsacales |
| *Lomelosia olgae* | Caprifoliaceae | Dipsacales |
| *Lotus herbaceus* | Fabaceae | Fabales |
| *Lunaria rediviva* | Brassicaceae | Brassicales |
| *Mahonia aquifolium* | Berberidaceae | Ranunculales |
| *Medicago falcata* | Fabaceae | Fabales |
| *Monarda fistulosa* | Lamiaceae | Lamiales |
| *Nicotiana tabacum* | Solanaceae | Solanales |
| *Papaver orientale* | Papaveraceae | Ranunculales |
| *Phlomis russeliana* | Lamiaceae | Lamiales |
| *Polygonatum biflorum* | Asparagaceae | Asparagales |
| *Potentilla kurdica* | Rosaceae | Rosales |
| *Potentilla recta* | Rosaceae | Rosales |
| *Pycnanthemum tenuifolium* | Lamiaceae | Lamiales |
| *Rhaponticoides ruthenica* | Asteraceae | Asterales |

| | | |
|---|---|---|
| *Rhododendron brachycarpum* | Ericaceae | Ericales |
| *Rhododendron degronianum* | Ericaceae | Ericales |
| *Rhododendron thomsonii* | Ericaceae | Ericales |
| *Rhododendron wadanum* | Ericaceae | Ericales |
| *Ribes sanguineum* | Grossulariaceae | Saxifragales |
| *Salvia ringens* | Lamiaceae | Lamiales |
| *Salvia verticillata* | Lamiaceae | Lamiales |
| *Saponaria sicula* | Caryophyllaceae | Caryophyllales |
| *Scabiosa triandra* | Caprifoliaceae | Dipsacales |
| *Scilla luciliae* | Asparagaceae | Asparagales |
| *Silene latifolia* | Caryophyllaceae | Caryophyllales |
| *Silene schafta* | Caryophyllaceae | Caryophyllales |
| *Silphium integrifolium* | Asteraceae | Asterales |
| *Solanum incanum* | Solanaceae | Solanales |
| *Succisa pratensis* | Caprifoliaceae | Dipsacales |
| *Syringa persica* | Oleaceae | Lamiales |
| *Tanacetum coccineum* | Asteraceae | Asterales |
| *Taraxacum sonchoides* | Asteraceae | Asterales |
| *Tetradium daniellii* | Rutaceae | Sapindales |
| *Teucrium marum* | Lamiaceae | Lamiales |
| *Thunbergia alata* | Acanthaceae | Lamiales |
| *Thymus praecox* | Lamiaceae | Lamiales |
| *Thymus pulegioides* | Lamiaceae | Lamiales |
| *Tropaeolum majus* | Tropaeolaceae | Brassicales |
| *Verbascum nigrum* | Scrophulariaceae | Lamiales |
| *Verbena hastata* | Verbenaceae | Lamiales |
| *Vernonia fasciculata* | Asteraceae | Asterales |
| *Veronica austriaca* | Plantaginaceae | Lamiales |
| *Veronica chamaedrys* | Plantaginaceae | Lamiales |
| *Veronica longifolia* | Plantaginaceae | Lamiales |
| *Veronica orchidea* | Plantaginaceae | Lamiales |
| *Vicia oroboides* | Fabaceae | Fabales |
| *Vicia tenuifolia* | Fabaceae | Fabales |
| *Viola striata* | Violaceae | Malpighiales |
| *Vitex negundo* | Lamiaceae | Lamiales |
| *Weigela florida* | Caprifoliaceae | Dipsacales |
| *Xanthoceras sorbifolium* | Poaceae | Poales |

**Table S5.** Non-focal plant species included in the study, with sampling time (minutes) per botanical garden (Leipzig, Halle, and Jena). The table also reports the total sampling time per species across all gardens. Species sampled in all three gardens are highlighted in bold. Total sampling time was 9 h in Leipzig, 10 h in Halle, and 10 h in Jena, summing to 29 h overall. Focal plant species are not included in this table, although they were occasionally recorded during random census sampling outside focal observation locations.

| Species | Leipzig (min) | Halle (min) | Jena (min) | Total time (min) |
|---|---|---|---|---|
| ***Achillea filipendulina*** | 6 | 3 | 9 | 18 |
| *Aconitum lycoctonum* | 0 | 3 | 0 | 3 |
| *Aegonychon purpurocaeruleum* | 0 | 6 | 0 | 6 |
| *Allium cernuum* | 0 | 6 | 3 | 9 |
| *Allium tuberosum* | 0 | 6 | 3 | 9 |
| *Allium turkestanicum* | 0 | 3 | 0 | 3 |
| *Alyssum saxatile* | 0 | 3 | 0 | 3 |
| *Anthriscus sylvestris* | 0 | 3 | 0 | 3 |
| *Arabis alpina* | 0 | 3 | 3 | 6 |
| *Asclepias syriaca* | 0 | 3 | 0 | 3 |
| *Asparagus oligoclonos* | 0 | 3 | 0 | 3 |
| *Astragalus alopecurus* | 0 | 3 | 0 | 3 |
| *Aubrieta deltoidea* | 0 | 6 | 3 | 9 |
| *Aubrieta olympica* | 0 | 3 | 0 | 3 |
| *Aubrieta parviflora* | 0 | 3 | 0 | 3 |
| *Aubrieta pinardii* | 0 | 3 | 24 | 27 |
| *Ballota nigra* | 0 | 3 | 0 | 3 |
| *Baptisia australis* | 0 | 6 | 6 | 12 |
| *Barbarea vulgaris* | 0 | 3 | 0 | 3 |
| ***Bellis perennis*** | 6 | 3 | 6 | 15 |
| *Bupleurum aureum* | 0 | 3 | 0 | 3 |
| *Campanula poscharskyana* | 0 | 3 | 0 | 3 |
| *Centaurea cyanus* | 0 | 3 | 0 | 3 |
| *Cephalaria uralensis* | 3 | 6 | 0 | 9 |
| *Cerinthe minor* | 0 | 6 | 0 | 6 |
| *Chrysojasminum fruticans* | 0 | 3 | 0 | 3 |
| *Cistus salviifolius* | 0 | 3 | 0 | 3 |
| *Clinopodium nepeta* | 0 | 6 | 0 | 6 |
| ***Corydalis cava*** | 3 | 34 | 11 | 48 |
| *Cosmos sulphureus* | 0 | 6 | 0 | 6 |
| *Cotoneaster adpressus* | 0 | 3 | 0 | 3 |
| *Crambe maritima* | 0 | 3 | 0 | 3 |
| *Cylindropuntia imbricata* | 0 | 3 | 0 | 3 |
| *Daucus carota* | 0 | 3 | 3 | 6 |
| *Delosperma cooperi* | 0 | 3 | 0 | 3 |
| *Delphinium elatum* | 0 | 3 | 0 | 3 |
| *Digitalis purpurea* | 0 | 3 | 0 | 3 |
| *Drymocallis rupestris* | 0 | 6 | 0 | 6 |
| *Erodium cicutarium* | 0 | 3 | 0 | 3 |
| *Eryngium caeruleum* | 0 | 3 | 0 | 3 |
| *Eryngium maritimum* | 0 | 3 | 0 | 3 |
| *Eryngium planum* | 0 | 3 | 0 | 3 |
| *Erysimum pulchellum* | 3 | 3 | 0 | 6 |
| *Eschscholzia californica* | 0 | 3 | 0 | 3 |

| | | | | |
|---|---|---|---|---|
| *Ferula assa-foetida* | 0 | 3 | 0 | 3 |
| *Foeniculum vulgare* | 0 | 3 | 0 | 3 |
| *Genista hispanica* | 0 | 6 | 0 | 6 |
| *Genista radiata* | 0 | 3 | 0 | 3 |
| *Geum ternatum* | 0 | 3 | 0 | 3 |
| *Globularia trichosantha* | 0 | 3 | 0 | 3 |
| *Gypsophila paniculata* | 0 | 3 | 0 | 3 |
| *Helianthemum apenninum* | 0 | 3 | 0 | 3 |
| *Heuchera grossulariifolia* | 0 | 3 | 0 | 3 |
| *Hippocrepis emerus* | 0 | 6 | 0 | 6 |
| *Hippomarathrum vulgare* | 0 | 3 | 0 | 3 |
| *Hypochaeris radicata* | 0 | 3 | 0 | 3 |
| *Impatiens glandulifera* | 0 | 3 | 0 | 3 |
| *Inula helenium* | 0 | 3 | 0 | 3 |
| *Ipheion uniflorum* | 0 | 1 | 18 | 19 |
| *Knautia dipsacifolia* | 3 | 6 | 0 | 9 |
| ***Lamium orvala*** | 9 | 3 | 15 | 27 |
| *Lamium purpureum* | 0 | 3 | 0 | 3 |
| *Liatris pycnostachya* | 0 | 3 | 0 | 3 |
| *Limnanthes douglasii* | 9 | 3 | 0 | 12 |
| *Limonium gmelini* | 0 | 3 | 3 | 6 |
| *Limonium sinuatum* | 0 | 3 | 0 | 3 |
| *Lythrum salicaria* | 3 | 18 | 0 | 21 |
| *Magnolia stellata* | 0 | 1 | 0 | 1 |
| *Malcolmia maritima* | 0 | 3 | 0 | 3 |
| *Malva thuringiaca* | 0 | 9 | 3 | 12 |
| *Marrubium peregrinum* | 0 | 9 | 0 | 9 |
| *Melilotus albus* | 0 | 6 | 0 | 6 |
| *Monarda citriodora* | 0 | 3 | 0 | 3 |
| *Moricandia arvensis* | 0 | 3 | 0 | 3 |
| *Muscari armeniacum* | 0 | 4 | 3 | 7 |
| ***Muscari botryoides*** | 27 | 30 | 18 | 75 |
| *Muscari comosum* | 0 | 3 | 0 | 3 |
| *Nepeta clarkei* | 0 | 6 | 0 | 6 |
| *Oenothera odorata* | 0 | 3 | 0 | 3 |
| *Ornithogalum nutans* | 0 | 6 | 3 | 9 |
| ***Paeonia daurica*** | 3 | 3 | 6 | 12 |
| *Penstemon hartwegii* | 0 | 3 | 0 | 3 |
| *Pentaglottis sempervirens* | 0 | 6 | 0 | 6 |
| *Phacelia congesta* | 0 | 3 | 0 | 3 |
| *Phedimus kamtschaticus* | 0 | 3 | 0 | 3 |
| *Phlomoides tuberosa* | 0 | 3 | 0 | 3 |
| *Pieris japonica* | 0 | 7 | 0 | 7 |
| *Potentilla delphinensis* | 0 | 3 | 0 | 3 |
| *Potentilla montana* | 0 | 3 | 0 | 3 |
| *Potentilla sterilis* | 0 | 1 | 0 | 1 |
| *Primula vulgaris* | 0 | 4 | 3 | 7 |
| *Prunella grandiflora* | 0 | 3 | 0 | 3 |
| *Prunus tenella* | 0 | 3 | 0 | 3 |
| *Pulmonaria mollis* | 0 | 37 | 0 | 37 |
| *Ranunculus bulbosus* | 0 | 9 | 0 | 9 |
| *Ranunculus caucasicus* | 0 | 3 | 0 | 3 |
| ***Ranunculus ficaria*** | 3 | 3 | 3 | 9 |
| *Ranunculus lanuginosus* | 0 | 3 | 0 | 3 |
| *Reseda alba* | 0 | 3 | 0 | 3 |
| *Reseda lutea* | 0 | 3 | 0 | 3 |

| | | | | |
|---|---|---|---|---|
| *Reseda odorata* | 0 | 3 | 0 | 3 |
| *Rhododendron smirnowii* | 0 | 3 | 0 | 3 |
| *Rosa foetida* | 0 | 3 | 0 | 3 |
| *Rudbeckia laciniata* | 0 | 3 | 0 | 3 |
| *Salvia sclarea* | 6 | 6 | 0 | 12 |
| *Satureja montana* | 0 | 9 | 0 | 9 |
| *Scutellaria baicalensis* | 0 | 3 | 0 | 3 |
| *Seseli gummiferum* | 0 | 3 | 0 | 3 |
| *Silene chungtienensis* | 0 | 3 | 0 | 3 |
| *Sisymbrium officinale* | 0 | 3 | 0 | 3 |
| *Stachys byzantina* | 9 | 3 | 0 | 12 |
| *Succisella inflexa* | 0 | 3 | 0 | 3 |
| ***Symphytum officinale*** | 12 | 3 | 3 | 18 |
| *Syringa tomentella* | 0 | 3 | 0 | 3 |
| *Tagetes erecta* | 0 | 3 | 0 | 3 |
| *Taraxacum officinale* | 6 | 9 | 0 | 15 |
| *Teucrium scorodonia* | 0 | 3 | 0 | 3 |
| *Thalictrum flavum* | 0 | 3 | 0 | 3 |
| *Thymus vulgaris* | 0 | 3 | 0 | 3 |
| *Trifolium rubens* | 0 | 3 | 3 | 6 |
| *Valeriana phu* | 0 | 3 | 0 | 3 |
| *Verbena rigida* | 0 | 6 | 0 | 6 |
| *Wisteria sinensis* | 0 | 3 | 0 | 3 |
| *Zinnia elegans* | 0 | 6 | 0 | 6 |
| *Acantholimon ulicinum* | 0 | 0 | 6 | 6 |
| *Agapanthus africanus* | 0 | 0 | 3 | 3 |
| *Alkanna orientalis* | 0 | 0 | 9 | 9 |
| *Anemone ranunculoides* | 0 | 0 | 3 | 3 |
| *Anthyllis vulneraria* | 0 | 0 | 9 | 9 |
| *Bergenia ciliata* | 0 | 0 | 3 | 3 |
| *Betonica alopecuros* | 0 | 0 | 3 | 3 |
| *Borago officinalis* | 0 | 0 | 3 | 3 |
| *Brassica oleracea* | 0 | 0 | 3 | 3 |
| *Buddleja davidii* | 0 | 0 | 3 | 3 |
| *Campanula alliariifolia* | 3 | 0 | 6 | 9 |
| *Campanula glomerata* | 0 | 0 | 3 | 3 |
| *Campanula lactiflora* | 0 | 0 | 6 | 6 |
| *Campanula persicifolia* | 0 | 0 | 3 | 3 |
| *Campanula pyramidalis* | 0 | 0 | 3 | 3 |
| *Campanula rapunculoides* | 0 | 0 | 6 | 6 |
| *Cardamine pratensis* | 0 | 0 | 21 | 21 |
| *Centaurea scabiosa* | 12 | 0 | 3 | 15 |
| *Centaurea stoebe* | 0 | 0 | 3 | 3 |
| *Centaurea subtilis* | 0 | 0 | 3 | 3 |
| *Cerastium arvense* | 0 | 0 | 6 | 6 |
| *Chamaecytisus austriacus* | 0 | 0 | 3 | 3 |
| *Chamaecytisus hirsutus* | 0 | 0 | 3 | 3 |
| *Cichorium intybus* | 3 | 0 | 3 | 6 |
| *Coronilla orientalis* | 0 | 0 | 3 | 3 |
| *Cytisophyllum sessilifolium* | 0 | 0 | 9 | 9 |
| *Dasiphora fruticosa* | 3 | 0 | 3 | 6 |
| *Desmodium canadense* | 0 | 0 | 3 | 3 |
| *Dianthus petraeus* | 0 | 0 | 3 | 3 |
| *Digitalis grandiflora* | 0 | 0 | 3 | 3 |
| *Digitalis obscura* | 0 | 0 | 3 | 3 |
| *Dracocephalum rupestre* | 0 | 0 | 3 | 3 |

| | | | | |
|---|---|---|---|---|
| *Erigeron annuus* | 0 | 0 | 6 | 6 |
| *Euphorbia nicaeensis* | 0 | 0 | 3 | 3 |
| *Eutrochium maculatum* | 0 | 0 | 3 | 3 |
| *Fagopyrum esculentum* | 0 | 0 | 3 | 3 |
| *Genista germanica* | 0 | 0 | 6 | 6 |
| *Geranium molle* | 0 | 0 | 3 | 3 |
| *Globularia cordifolia* | 0 | 0 | 3 | 3 |
| *Helichrysum arenarium* | 0 | 0 | 3 | 3 |
| *Heracleum sphondylium* | 0 | 0 | 3 | 3 |
| *Heuchera villosa* | 0 | 0 | 3 | 3 |
| *Hypericum fragile* | 0 | 0 | 3 | 3 |
| *Hypericum hookerianum* | 0 | 0 | 6 | 6 |
| *Inula salicina* | 0 | 0 | 3 | 3 |
| *Iris germanica* | 0 | 0 | 3 | 3 |
| *Iris pseudacorus* | 0 | 0 | 3 | 3 |
| *Jacobaea erucifolia* | 0 | 0 | 3 | 3 |
| *Knautia arvensis* | 9 | 0 | 3 | 12 |
| *Lactuca macrophylla* | 0 | 0 | 3 | 3 |
| *Lactuca plumieri* | 0 | 0 | 3 | 3 |
| *Lamium galeobdolon* | 0 | 0 | 9 | 9 |
| *Lamprocapnos spectabilis* | 0 | 0 | 3 | 3 |
| *Lathyrus aureus* | 0 | 0 | 6 | 6 |
| *Lathyrus niger* | 0 | 0 | 3 | 3 |
| *Lathyrus pratensis* | 0 | 0 | 3 | 3 |
| *Lilium martagon* | 0 | 0 | 3 | 3 |
| *Lindelofia anchusoides* | 0 | 0 | 3 | 3 |
| *Lomelosia albocincta* | 0 | 0 | 3 | 3 |
| *Lonicera floribunda* | 0 | 0 | 6 | 6 |
| *Lysimachia vulgaris* | 0 | 0 | 6 | 6 |
| *Medicago sativa* | 0 | 0 | 6 | 6 |
| *Omphalodes verna* | 0 | 0 | 15 | 15 |
| *Paeonia mascula* | 0 | 0 | 15 | 15 |
| *Papaver nudicaule* | 0 | 0 | 6 | 6 |
| *Papaver somniferum* | 0 | 0 | 3 | 3 |
| *Pentanema helenioides* | 0 | 0 | 3 | 3 |
| *Phacelia tanacetifolia* | 0 | 0 | 3 | 3 |
| *Phedimus aizoon* | 3 | 0 | 3 | 6 |
| *Philadelphus pekinensis* | 0 | 0 | 9 | 9 |
| *Phlox paniculata* | 0 | 0 | 3 | 3 |
| *Pilosella officinarum* | 0 | 0 | 3 | 3 |
| *Rhododendron saluenense* | 0 | 0 | 3 | 3 |
| *Rosa spinosissima* | 0 | 0 | 3 | 3 |
| *Rudbeckia fulgida* | 0 | 0 | 3 | 3 |
| *Salvia nutans* | 0 | 0 | 3 | 3 |
| *Satureja parnassica* | 0 | 0 | 3 | 3 |
| *Scabiosa canescens* | 0 | 0 | 3 | 3 |
| *Scrophularia versicolor* | 0 | 0 | 3 | 3 |
| *Scutellaria altissima* | 0 | 0 | 3 | 3 |
| *Scutellaria cordifolia* | 0 | 0 | 6 | 6 |
| *Seseli montanum* | 3 | 0 | 3 | 6 |
| *Solidago flexicaulis* | 0 | 0 | 6 | 6 |
| *Spartium junceum* | 0 | 0 | 6 | 6 |
| *Spiraea chamaedryfolia* | 0 | 0 | 3 | 3 |
| *Stellaria palustris* | 0 | 0 | 18 | 18 |
| *Symphytum bulbosum* | 6 | 0 | 3 | 9 |
| *Teucrium canadense* | 0 | 0 | 3 | 3 |

| | | | | |
|---|---|---|---|---|
| *Teucrium chamaedrys* | 0 | 0 | 3 | 3 |
| *Tilia henryana* | 0 | 0 | 3 | 3 |
| *Trifolium pratense* | 0 | 0 | 21 | 21 |
| *Tripleurospermum maritimum* | 0 | 0 | 3 | 3 |
| *Trommsdorffia maculata* | 3 | 0 | 3 | 6 |
| *Verbascum phoeniceum* | 0 | 0 | 3 | 3 |
| *Vicia sepium* | 0 | 0 | 6 | 6 |
| *Aethionema grandiflorum* | 3 | 0 | 0 | 3 |
| *Ajuga reptans* | 3 | 0 | 0 | 3 |
| *Alisma plantago-aquatica* | 6 | 0 | 0 | 6 |
| *Allium senescens* | 3 | 0 | 0 | 3 |
| *Anaphalis margaritacea* | 3 | 0 | 0 | 3 |
| *Anchusa officinalis* | 12 | 0 | 0 | 12 |
| *Aster mongolicus* | 3 | 0 | 0 | 3 |
| *Aurinia saxatilis* | 3 | 0 | 0 | 3 |
| *Carlina vulgaris* | 3 | 0 | 0 | 3 |
| *Centaurea nigra* | 3 | 0 | 0 | 3 |
| *Centaurea sadleriana* | 3 | 0 | 0 | 3 |
| *Cephalaria gigantea* | 3 | 0 | 0 | 3 |
| *Cephalaria procera* | 3 | 0 | 0 | 3 |
| *Chaenorhinum origanifolium* | 3 | 0 | 0 | 3 |
| *Cirsium vulgare* | 3 | 0 | 0 | 3 |
| *Clinopodium album* | 3 | 0 | 0 | 3 |
| *Convolvulus tricolor* | 3 | 0 | 0 | 3 |
| *Coristospermum lucidum* | 3 | 0 | 0 | 3 |
| *Cuphea viscosissima* | 9 | 0 | 0 | 9 |
| *Dasiphora davurica* | 9 | 0 | 0 | 9 |
| *Echinacea angustifolia* | 6 | 0 | 0 | 6 |
| *Echinops pungens* | 3 | 0 | 0 | 3 |
| *Echium vulgare* | 3 | 0 | 0 | 3 |
| *Erica carnea* | 6 | 0 | 0 | 6 |
| *Eriocapitella tomentosa* | 3 | 0 | 0 | 3 |
| *Erodium manescavi* | 3 | 0 | 0 | 3 |
| *Euphorbia characias* | 6 | 0 | 0 | 6 |
| *Euphorbia seguieriana* | 9 | 0 | 0 | 9 |
| *Ferulago aucheri* | 6 | 0 | 0 | 6 |
| *Forsythia intermedia* | 3 | 0 | 0 | 3 |
| *Geranium macrorrhizum* | 9 | 0 | 0 | 9 |
| *Helichrysum italicum* | 3 | 0 | 0 | 3 |
| *Hieracium murorum* | 3 | 0 | 0 | 3 |
| *Impatiens balsamina* | 3 | 0 | 0 | 3 |
| *Inula oculus-christi* | 3 | 0 | 0 | 3 |
| *Klasea radiata* | 3 | 0 | 0 | 3 |
| *Knautia macedonica* | 6 | 0 | 0 | 6 |
| *Lamium flexuosum* | 3 | 0 | 0 | 3 |
| *Lamium hybridum* | 10 | 0 | 0 | 10 |
| *Lathyrus roseus* | 3 | 0 | 0 | 3 |
| *Lomelosia cretica* | 6 | 0 | 0 | 6 |
| *Lomelosia olgae* | 6 | 0 | 0 | 6 |
| *Lotus herbaceus* | 3 | 0 | 0 | 3 |
| *Lunaria rediviva* | 3 | 0 | 0 | 3 |
| *Mahonia aquifolium* | 6 | 0 | 0 | 6 |
| *Medicago falcata* | 3 | 0 | 0 | 3 |
| *Monarda fistulosa* | 3 | 0 | 0 | 3 |
| *Nicotiana tabacum* | 3 | 0 | 0 | 3 |
| *Papaver orientale* | 3 | 0 | 0 | 3 |

| | | | | |
|---|---|---|---|---|
| *Phlomis russeliana* | 3 | 0 | 0 | 3 |
| *Polygonatum biflorum* | 6 | 0 | 0 | 6 |
| *Potentilla kurdica* | 9 | 0 | 0 | 9 |
| *Potentilla recta* | 3 | 0 | 0 | 3 |
| *Pycnanthemum tenuifolium* | 6 | 0 | 0 | 6 |
| *Rhaponticoides ruthenica* | 3 | 0 | 0 | 3 |
| *Rhododendron brachycarpum* | 6 | 0 | 0 | 6 |
| *Rhododendron degronianum* | 3 | 0 | 0 | 3 |
| *Rhododendron thomsonii* | 3 | 0 | 0 | 3 |
| *Rhododendron wadanum* | 6 | 0 | 0 | 6 |
| *Ribes sanguineum* | 15 | 0 | 0 | 15 |
| *Salvia ringens* | 3 | 0 | 0 | 3 |
| *Salvia verticillata* | 6 | 0 | 0 | 6 |
| *Saponaria sicula* | 6 | 0 | 0 | 6 |
| *Scabiosa triandra* | 3 | 0 | 0 | 3 |
| *Scilla luciliae* | 15 | 0 | 0 | 15 |
| *Silene latifolia* | 3 | 0 | 0 | 3 |
| *Silene schafta* | 3 | 0 | 0 | 3 |
| *Silphium integrifolium* | 6 | 0 | 0 | 6 |
| *Solanum incanum* | 3 | 0 | 0 | 3 |
| *Succisa pratensis* | 6 | 0 | 0 | 6 |
| *Syringa persica* | 3 | 0 | 0 | 3 |
| *Tanacetum coccineum* | 3 | 0 | 0 | 3 |
| *Taraxacum sonchoides* | 3 | 0 | 0 | 3 |
| *Tetradium daniellii* | 3 | 0 | 0 | 3 |
| *Teucrium marum* | 3 | 0 | 0 | 3 |
| *Thunbergia alata* | 3 | 0 | 0 | 3 |
| *Thymus praecox* | 3 | 0 | 0 | 3 |
| *Thymus pulegioides* | 3 | 0 | 0 | 3 |
| *Tropaeolum majus* | 3 | 0 | 0 | 3 |
| *Verbascum nigrum* | 3 | 0 | 0 | 3 |
| *Verbena hastata* | 3 | 0 | 0 | 3 |
| *Vernonia fasciculata* | 3 | 0 | 0 | 3 |
| *Veronica austriaca* | 3 | 0 | 0 | 3 |
| *Veronica chamaedrys* | 3 | 0 | 0 | 3 |
| *Veronica longifolia* | 3 | 0 | 0 | 3 |
| *Veronica orchidea* | 3 | 0 | 0 | 3 |
| *Vicia oroboides* | 3 | 0 | 0 | 3 |
| *Vicia tenuifolia* | 6 | 0 | 0 | 6 |
| *Viola striata* | 9 | 0 | 0 | 9 |
| *Vitex negundo* | 12 | 0 | 0 | 12 |
| *Weigela florida* | 3 | 0 | 0 | 3 |
| *Xanthoceras sorbifolium* | 5 | 0 | 0 | 5 |

**Table S6.** Pollinator species recorded in the study, including their family and order. The table also indicates the botanical gardens in which each species was observed, using abbreviated codes: H = Halle, J = Jena, and L = Leipzig.

| Species | Family | Order | Garden |
|---|---|---|---|
| *Andrena bicolor* | Andrenidae | Hymenoptera | HL |
| *Andrena chrysopus* | Andrenidae | Hymenoptera | L |
| *Andrena curvungula* | Andrenidae | Hymenoptera | L |
| *Andrena flavipes* | Andrenidae | Hymenoptera | HJL |
| *Andrena fulvago* | Andrenidae | Hymenoptera | L |
| *Andrena gelriae* | Andrenidae | Hymenoptera | J |
| *Andrena gravida* | Andrenidae | Hymenoptera | L |
| *Andrena hattorfiana* | Andrenidae | Hymenoptera | HL |
| *Andrena labiata* | Andrenidae | Hymenoptera | L |
| *Andrena minutula* | Andrenidae | Hymenoptera | HJL |
| *Andrena minutuloides* | Andrenidae | Hymenoptera | HL |
| *Andrena nigroaenea* | Andrenidae | Hymenoptera | L |
| *Andrena nitida* | Andrenidae | Hymenoptera | J |
| *Andrena pilipes* | Andrenidae | Hymenoptera | HL |
| *Andrena strohmella* | Andrenidae | Hymenoptera | H |
| *Anthophora aestivalis* | Apidae | Hymenoptera | HJL |
| *Anthophora furcata* | Apidae | Hymenoptera | H |
| *Anthophora plumipes* | Apidae | Hymenoptera | HJL |
| *Anthophora quadrimaculata* | Apidae | Hymenoptera | HJL |
| *Apis mellifera* | Apidae | Hymenoptera | HJL |
| *Bombus campestris* | Apidae | Hymenoptera | L |
| *Bombus cryptarum* | Apidae | Hymenoptera | H |
| *Bombus hortorum* | Apidae | Hymenoptera | HJL |
| *Bombus hypnorum* | Apidae | Hymenoptera | HJL |
| *Bombus lapidarius* | Apidae | Hymenoptera | HJL |
| *Bombus lucorum* | Apidae | Hymenoptera | HJ |
| *Bombus pascuorum* | Apidae | Hymenoptera | HJL |
| *Bombus pratorum* | Apidae | Hymenoptera | HJL |
| *Bombus rupestris* | Apidae | Hymenoptera | H |
| *Bombus semenoviellus* | Apidae | Hymenoptera | H |
| *Bombus soroeensis* | Apidae | Hymenoptera | J |
| *Bombus subterraneus* | Apidae | Hymenoptera | H |
| *Bombus terrestris* | Apidae | Hymenoptera | HJL |
| *Bombus vestalis* | Apidae | Hymenoptera | HJL |
| *Ceratina cyanea* | Apidae | Hymenoptera | L |
| *Eucera nigrescens* | Apidae | Hymenoptera | HJL |
| *Melecta albifrons* | Apidae | Hymenoptera | HJ |
| *Nomada bifasciata* | Apidae | Hymenoptera | H |
| *Nomada fabriciana* | Apidae | Hymenoptera | L |
| *Nomada flavoguttata* | Apidae | Hymenoptera | HL |
| *Nomada fucata* | Apidae | Hymenoptera | H |
| *Nomada fulvicornis* | Apidae | Hymenoptera | H |
| *Nomada lathburiana* | Apidae | Hymenoptera | H |
| *Nomada zonata* | Apidae | Hymenoptera | J |
| *Thyreus orbatus* | Apidae | Hymenoptera | HJ |
| *Xylocopa violacea* | Apidae | Hymenoptera | HJL |
| *Hedychrum gerstaeckeri* | Chrysididae | Hymenoptera | HL |
| *Hedychrum nobile* | Chrysididae | Hymenoptera | H |
| *Holopyga ignicollis* | Chrysididae | Hymenoptera | L |
| *Colletes daviesanus* | Colletidae | Hymenoptera | HJL |

| | | | |
|---|---|---|---|
| *Colletes hederae* | Colletidae | Hymenoptera | H |
| *Hylaeus brevicornis* | Colletidae | Hymenoptera | L |
| *Hylaeus communis* | Colletidae | Hymenoptera | HJL |
| *Hylaeus confusus* | Colletidae | Hymenoptera | L |
| *Hylaeus gredleri* | Colletidae | Hymenoptera | HL |
| *Hylaeus hyalinatus* | Colletidae | Hymenoptera | L |
| *Hylaeus nigritus* | Colletidae | Hymenoptera | HJL |
| *Hylaeus punctatus* | Colletidae | Hymenoptera | HJL |
| *Hylaeus signatus* | Colletidae | Hymenoptera | H |
| *Hylaeus sinuatus* | Colletidae | Hymenoptera | J |
| *Hylaeus styriacus* | Colletidae | Hymenoptera | L |
| *Cerceris quadricincta* | Crabronidae | Hymenoptera | J |
| *Cerceris quinquefasciata* | Crabronidae | Hymenoptera | L |
| *Cerceris rybyensis* | Crabronidae | Hymenoptera | HJL |
| *Crossocerus podagricus* | Crabronidae | Hymenoptera | H |
| *Dinetus pictus* | Crabronidae | Hymenoptera | HL |
| *Ectemnius lituratus* | Crabronidae | Hymenoptera | J |
| *Nysson maculosus* | Crabronidae | Hymenoptera | L |
| *Philanthus triangulum* | Crabronidae | Hymenoptera | HL |
| *Gymnomerus laevipes* | Eumenidae | Hymenoptera | JL |
| *Polistes dominula* | Eumenidae | Hymenoptera | HJL |
| *Halictus rubicundus* | Halictidae | Hymenoptera | H |
| *Halictus scabiosae* | Halictidae | Hymenoptera | HJL |
| *Halictus subauratus* | Halictidae | Hymenoptera | HJL |
| *Halictus tumulorum* | Halictidae | Hymenoptera | HJL |
| *Lasioglossum calceatum* | Halictidae | Hymenoptera | HJL |
| *Lasioglossum fulvicorne* | Halictidae | Hymenoptera | H |
| *Lasioglossum interruptum* | Halictidae | Hymenoptera | J |
| *Lasioglossum laticeps* | Halictidae | Hymenoptera | HL |
| *Lasioglossum leucozonium* | Halictidae | Hymenoptera | HL |
| *Lasioglossum minutissimum* | Halictidae | Hymenoptera | L |
| *Lasioglossum morio* | Halictidae | Hymenoptera | HJL |
| *Lasioglossum nitidiusculum* | Halictidae | Hymenoptera | H |
| *Lasioglossum nitidulum* | Halictidae | Hymenoptera | H |
| *Lasioglossum pallens* | Halictidae | Hymenoptera | JL |
| *Lasioglossum pauxillum* | Halictidae | Hymenoptera | HJL |
| *Lasioglossum politum* | Halictidae | Hymenoptera | HJL |
| *Lasioglossum pygmaeum* | Halictidae | Hymenoptera | L |
| *Lasioglossum villosulum* | Halictidae | Hymenoptera | HL |
| *Sphecodes albilabris* | Halictidae | Hymenoptera | HL |
| *Sphecodes ferruginatus* | Halictidae | Hymenoptera | H |
| *Sphecodes monilicornis* | Halictidae | Hymenoptera | H |
| *Sphecodes pellucidus* | Halictidae | Hymenoptera | HL |
| *Sphecodes puncticeps* | Halictidae | Hymenoptera | HL |
| *Anthidium manicatum* | Megachilidae | Hymenoptera | HJL |
| *Anthidium oblongatum* | Megachilidae | Hymenoptera | HL |
| *Chelostoma florisomne* | Megachilidae | Hymenoptera | HJ |
| *Chelostoma rapunculi* | Megachilidae | Hymenoptera | HJL |
| *Coelioxys afra* | Megachilidae | Hymenoptera | H |
| *Coelioxys aurolimbatus* | Megachilidae | Hymenoptera | H |
| *Coelioxys elongatus* | Megachilidae | Hymenoptera | HL |
| *Coelioxys rufescens* | Megachilidae | Hymenoptera | H |
| *Heriades truncorum* | Megachilidae | Hymenoptera | HJL |
| *Hoplitis adunca* | Megachilidae | Hymenoptera | HJL |
| *Megachile alpicola* | Megachilidae | Hymenoptera | J |
| *Megachile centuncularis* | Megachilidae | Hymenoptera | HJ |

| | | | |
|---|---|---|---|
| *Megachile circumcincta* | Megachilidae | Hymenoptera | JL |
| *Megachile ericetorum* | Megachilidae | Hymenoptera | HJL |
| *Megachile lagopoda* | Megachilidae | Hymenoptera | HJL |
| *Megachile maritima* | Megachilidae | Hymenoptera | HJL |
| *Megachile pilidens* | Megachilidae | Hymenoptera | HJ |
| *Megachile rotundata* | Megachilidae | Hymenoptera | HJL |
| *Megachile versicolor* | Megachilidae | Hymenoptera | J |
| *Megachile willughbiella* | Megachilidae | Hymenoptera | HJL |
| *Osmia aurulenta* | Megachilidae | Hymenoptera | HJL |
| *Osmia bicolor* | Megachilidae | Hymenoptera | H |
| *Osmia bicornis* | Megachilidae | Hymenoptera | HJL |
| *Osmia brevicornis* | Megachilidae | Hymenoptera | HJL |
| *Osmia caerulescens* | Megachilidae | Hymenoptera | HJL |
| *Osmia cornuta* | Megachilidae | Hymenoptera | HJL |
| *Pseudoanthidium nanum* | Megachilidae | Hymenoptera | L |
| *Stelis punctulatissima* | Megachilidae | Hymenoptera | JL |
| *Macropis fulvipes* | Melittidae | Hymenoptera | J |
| *Melitta haemorrhoidalis* | Melittidae | Hymenoptera | L |
| *Melitta nigricans* | Melittidae | Hymenoptera | H |
| *Sapygina decemguttata* | Sapygidae | Hymenoptera | H |
| *Scolia hirta* | Scoliidae | Hymenoptera | H |
| *Ammophila sabulosa* | Sphecidae | Hymenoptera | H |
| *Sphex funerarius* | Sphecidae | Hymenoptera | HJL |
| *Tiphia femorata* | Tiphiidae | Hymenoptera | HL |
| *Ancistrocerus claripennis* | Vespidae | Hymenoptera | H |
| *Ancistrocerus gazella* | Vespidae | Hymenoptera | H |
| *Ancistrocerus nigricornis* | Vespidae | Hymenoptera | H |
| *Dolichovespula saxonica* | Vespidae | Hymenoptera | H |
| *Vespa crabro* | Vespidae | Hymenoptera | HJ |
| *Vespula germanica* | Vespidae | Hymenoptera | HJ |
| *Anthomyia procellaris* | Anthomyiidae | Diptera | H |
| *Botanophila depressa* | Anthomyiidae | Diptera | L |
| *Botanophila striolata* | Anthomyiidae | Diptera | JL |
| *Leucophora personata* | Anthomyiidae | Diptera | J |
| *Bombylius discolor* | Bombyliidae | Diptera | HJL |
| *Bombylius major* | Bombyliidae | Diptera | HJL |
| *Bombylius venosus* | Bombyliidae | Diptera | HJL |
| *Hemipenthes maura* | Bombyliidae | Diptera | J |
| *Hemipenthes morio* | Bombyliidae | Diptera | H |
| *Villa modesta* | Bombyliidae | Diptera | L |
| *Bellardia viarum* | Calliphoridae | Diptera | H |
| *Bellardia vulgaris* | Calliphoridae | Diptera | HL |
| *Calliphora vicina* | Calliphoridae | Diptera | L |
| *Eurychaeta palpalis* | Calliphoridae | Diptera | HJL |
| *Lucilia sericata* | Calliphoridae | Diptera | HJL |
| *Dalmannia marginata* | Conopidae | Diptera | L |
| *Merziella longirostris* | Conopidae | Diptera | H |
| *Myopa testacea* | Conopidae | Diptera | L |
| *Physocephala rufipes* | Conopidae | Diptera | HL |
| *Sicus ferrugineus* | Conopidae | Diptera | HJL |
| *Minettia longipennis* | Lauxaniidae | Diptera | J |
| *Graphomya maculata* | Muscidae | Diptera | H |
| *Platystoma seminationis* | Platystomatidae | Diptera | L |
| *Pollenia amentaria* | Polleniidae | Diptera | H |
| *Pollenia pediculata* | Polleniidae | Diptera | H |
| *Pollenia vagabunda* | Polleniidae | Diptera | H |

| | | | |
|---|---|---|---|
| *Miltogramma germari* | Sarcophagidae | Diptera | H |
| *Miltogramma punctata* | Sarcophagidae | Diptera | HL |
| *Sarcophaga haemorrhoa* | Sarcophagidae | Diptera | J |
| *Sarcophaga incisilobata* | Sarcophagidae | Diptera | HJ |
| *Sarcophaga subvicina* | Sarcophagidae | Diptera | H |
| *Cheilosia aerea* | Syrphidae | Diptera | L |
| *Cheilosia pagana* | Syrphidae | Diptera | J |
| *Cheilosia ruffipes* | Syrphidae | Diptera | HJL |
| *Dasysyrphus albostriatus* | Syrphidae | Diptera | J |
| *Episyrphus balteatus* | Syrphidae | Diptera | HJL |
| *Eristalinus aeneus* | Syrphidae | Diptera | H |
| *Eristalis pertinax* | Syrphidae | Diptera | HJ |
| *Eristalis tenax* | Syrphidae | Diptera | HJL |
| *Eupeodes corollae* | Syrphidae | Diptera | HJL |
| *Lapposyrphus lapponicus* | Syrphidae | Diptera | L |
| *Meliscaeva auricollis* | Syrphidae | Diptera | J |
| *Merodon equestris* | Syrphidae | Diptera | L |
| *Myathropa florea* | Syrphidae | Diptera | HJL |
| *Paragus constrictus* | Syrphidae | Diptera | J |
| *Scaeva pyrastri* | Syrphidae | Diptera | HJ |
| *Sphaerophoria rueppellii* | Syrphidae | Diptera | J |
| *Sphaerophoria scripta* | Syrphidae | Diptera | HJL |
| *Syritta pipiens* | Syrphidae | Diptera | HJ |
| *Syrphus vitripennis* | Syrphidae | Diptera | HJ |
| *Trichopsomyia flavitarsis* | Syrphidae | Diptera | H |
| *Volucella inanis* | Syrphidae | Diptera | J |
| *Volucella pellucens* | Syrphidae | Diptera | J |
| *Volucella zonaria* | Syrphidae | Diptera | JL |
| *Cylindromyia brevicornis* | Tachinidae | Diptera | HJL |
| *Entomophaga nigrohalterata* | Tachinidae | Diptera | HJ |
| *Gymnosoma clavatum* | Tachinidae | Diptera | HJ |
| *Gymnosoma nudifrons* | Tachinidae | Diptera | JL |
| *Hebia flavipes* | Tachinidae | Diptera | J |
| *Litophasia hyalipennis* | Tachinidae | Diptera | H |
| *Microphthalma europaea* | Tachinidae | Diptera | L |
| *Nowickia ferox* | Tachinidae | Diptera | H |
| *Phasia barbifrons* | Tachinidae | Diptera | J |
| *Phasia pusilla* | Tachinidae | Diptera | J |
| *Prosena siberita* | Tachinidae | Diptera | H |
| *Tachina fera* | Tachinidae | Diptera | H |
| *Tachina nupta* | Tachinidae | Diptera | HL |
| *Anthaxia nitidula* | Buprestidae | Coleoptera | H |
| *Pseudovadonia livida* | Cerambycidae | Coleoptera | HL |
| *Stenurella melanura* | Cerambycidae | Coleoptera | H |
| *Timarcha goettingensis* | Chrysomelidae | Coleoptera | J |
| *Trichodes apiarius* | Cleridae | Coleoptera | J |
| *Coccinella septempunctata* | Coccinellidae | Coleoptera | HJL |
| *Harmonia axyridis* | Coccinellidae | Coleoptera | HJ |
| *Anthrenus scrophulariae* | Dermestidae | Coleoptera | H |
| *Anthrenus verbasci* | Dermestidae | Coleoptera | HJ |
| *Malachius bipustulatus* | Melyridae | Coleoptera | H |
| *Oedemera femorata* | Oedemeridae | Coleoptera | H |
| *Oedemera lurida* | Oedemeridae | Coleoptera | HJL |
| *Oedemera podagrariae* | Oedemeridae | Coleoptera | H |
| *Cetonia aurata* | Scarabaeidae | Coleoptera | HJL |
| *Oxythyrea funesta* | Scarabaeidae | Coleoptera | HJL |

| | | | |
|---|---|---|---|
| *Trichius gallicus* | Scarabaeidae | Coleoptera | HL |
| *Tropinota hirta* | Scarabaeidae | Coleoptera | H |
| *Valgus hemipterus* | Scarabaeidae | Coleoptera | HL |
| *Cydalima perspectalis* | Crambidae | Lepidoptera | L |
| *Aricia agestis* | Lycaenidae | Lepidoptera | J |
| *Polyommatus icarus* | Lycaenidae | Lepidoptera | HJL |
| *Autographa gamma* | Noctuidae | Lepidoptera | J |
| *Aglais io* | Nymphalidae | Lepidoptera | HJL |
| *Aglais urticae* | Nymphalidae | Lepidoptera | H |
| *Argynnis paphia* | Nymphalidae | Lepidoptera | J |
| *Dryas iulia* | Nymphalidae | Lepidoptera | L |
| *Issoria lathonia* | Nymphalidae | Lepidoptera | J |
| *Maniola jurtina* | Nymphalidae | Lepidoptera | HJ |
| *Polygonia c-album* | Nymphalidae | Lepidoptera | H |
| *Anthocharis cardamines* | Pieridae | Lepidoptera | HJL |
| *Gonepteryx rhamni* | Pieridae | Lepidoptera | H |
| *Pieris brassicae* | Pieridae | Lepidoptera | J |
| *Pieris napi* | Pieridae | Lepidoptera | HJL |
| *Pieris rapae* | Pieridae | Lepidoptera | HJL |
| *Macroglossum stellatarum* | Sphingidae | Lepidoptera | HJ |
| *Lygaeus equestris* | Lygaeidae | Hemiptera | L |
| *Carpocoris fuscispinus* | Pentatomidae | Hemiptera | J |
| *Graphosoma italicum* | Pentatomidae | Hemiptera | L |
| *Pyrrhocoris apterus* | Pyrrhocoridae | Hemiptera | H |
| *Pyrrhocoris marginatus* | Pyrrhocoridae | Hemiptera | J |

**Table S7.** Summary statistics (mean, standard deviation, minimum, and maximum) for plant and pollinator traits used in the analyses.

| Group | Traits | Mean | SD | Min | Max |
|---|---|---|---|---|---|
| a) Plant traits | Autonomous selfing | 6.5 | 21.7 | 0 | 88 |
| a) Plant traits | Corolla diameter | 26.2 | 22.1 | 4.5 | 115 |
| a) Plant traits | Flowers per plant | 551 | 1406.1 | 1 | 8000 |
| a) Plant traits | Nectar volume | 1.3 | 3.8 | 0 | 23.3 |
| a) Plant traits | Ovule number | 66.7 | 92 | 1 | 390 |
| a) Plant traits | Plant height | 0.5 | 0.33 | 0.04 | 1.49 |
| a) Plant traits | Pollen per flower | 381413.8 | 1461347.9 | 73.5 | 11994432 |
| a) Plant traits | Pollen size | 42.1 | 28.8 | 17.5 | 240 |
| a) Plant traits | Style length | 9.5 | 13.3 | 0 | 84.9 |
| b) Pollinator traits | Body length | 9.0 | 3.6 | 02.07 | 24 |
| b) Pollinator traits | IT | 3.1 | 1.4 | 1.18 | 9.1 |
| b) Pollinator traits | Proboscis length | 3.9 | 3.1 | 0.4 | 17.9 |

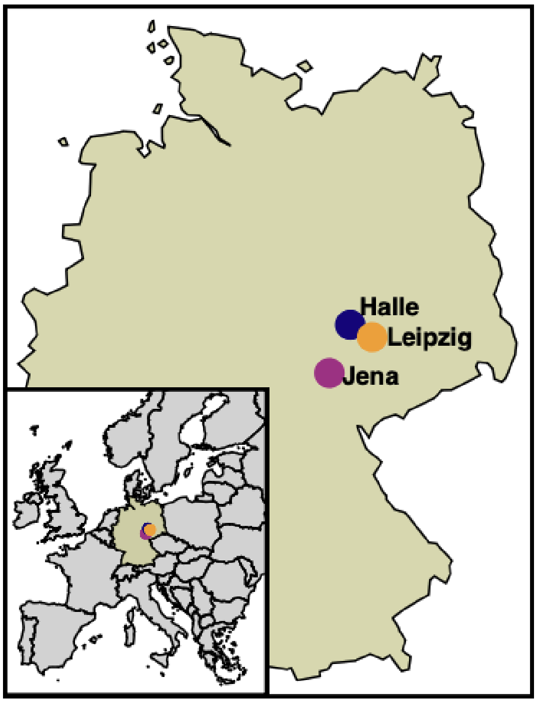


**Figure S1.** Map showing the locations of the three botanical gardens included in the study.

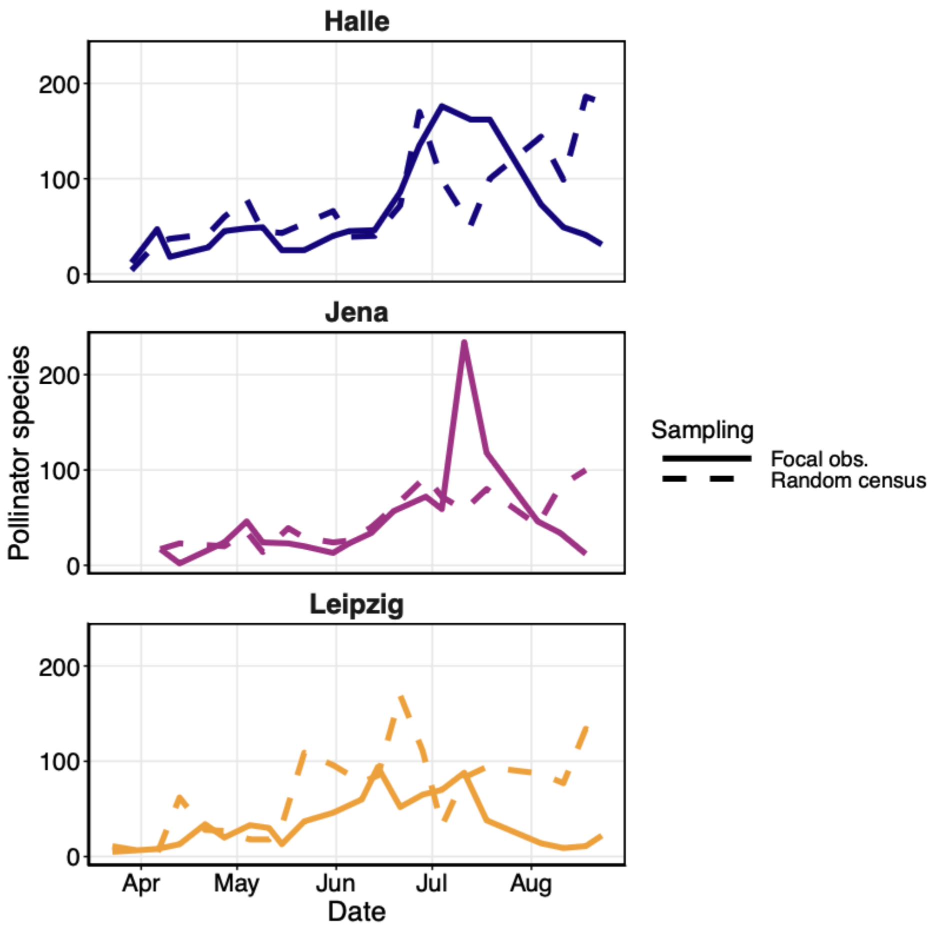


**Figure S2.** Temporal dynamics of plant-pollinator interactions across three botanical gardens. Solid lines represent focal observations on PhenObs plants, whereas dashed lines indicate random census observations across the gardens. Each panel corresponds to a different botanical garden: (a) Halle, (b) Leipzig, and (c) Jena.

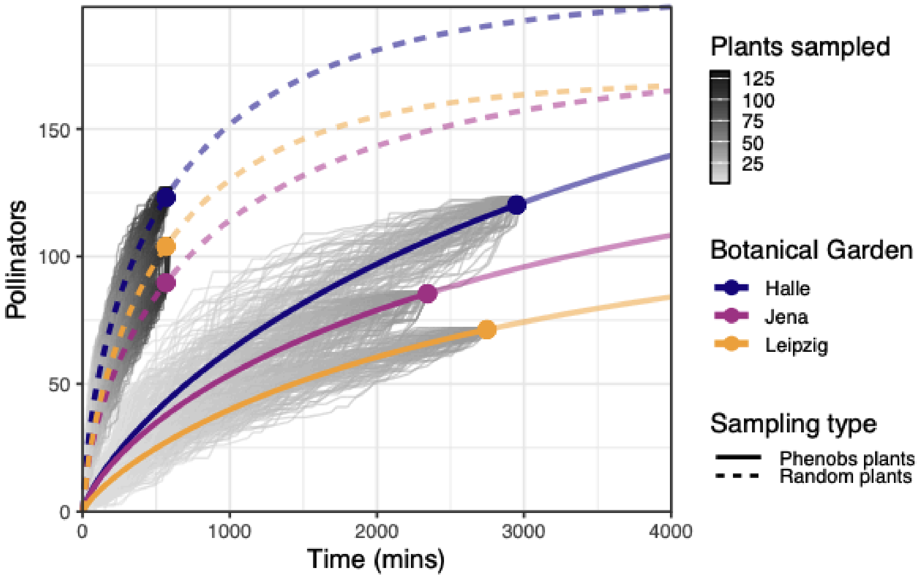


**Figure S3.** Accumulation curves of pollinator species across botanical gardens and sampling methods (dashed and solid lines). Each grey line represents one iteration of the accumulation curve, with a total of 100 iterations per garden and sampling method. The coloured lines represent the rarefied mean accumulation curves, with the dot indicating the total number of observed species. The segments of the lines extending beyond the dot represent the extrapolated number of expected species.

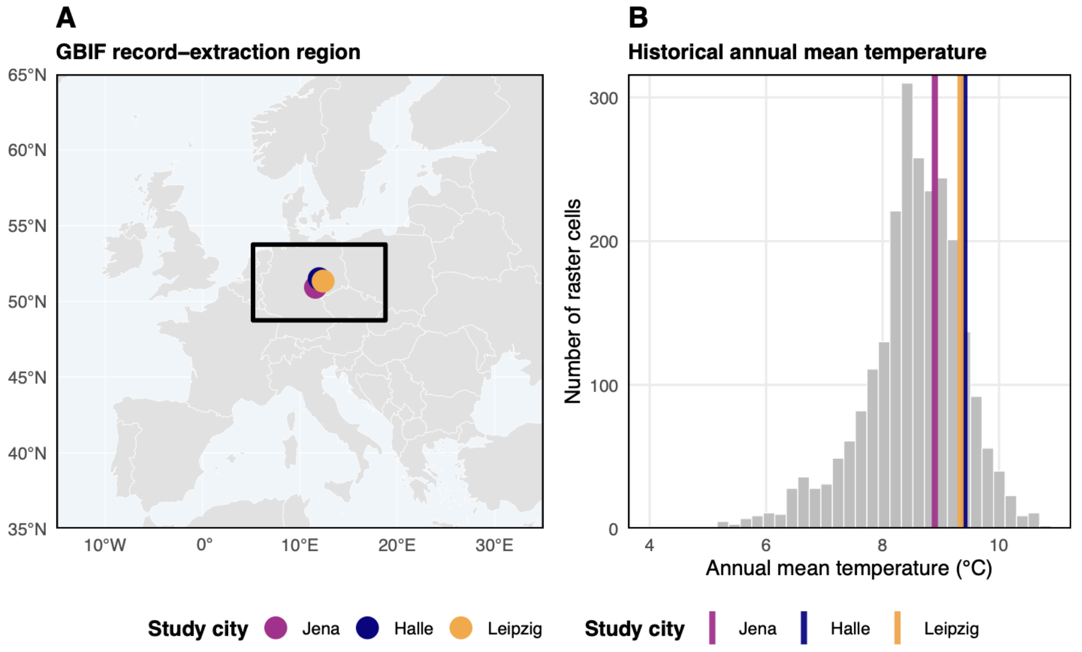


**Figure S4.** Spatial and climatic characterization of the GBIF record-extraction region used to estimate pollinator phenology. **(A)** Geographic extent (black rectangle) centred on Jena, Halle, and Leipzig. **(B)** Distribution of long-term annual mean temperature (WorldClim v2.1 climatological normals) within the extraction region. Colored vertical lines denote the temperatures of the three study cities, supporting the use of the selected region as a compromise between maximizing occurrence records and limiting climatic heterogeneity.

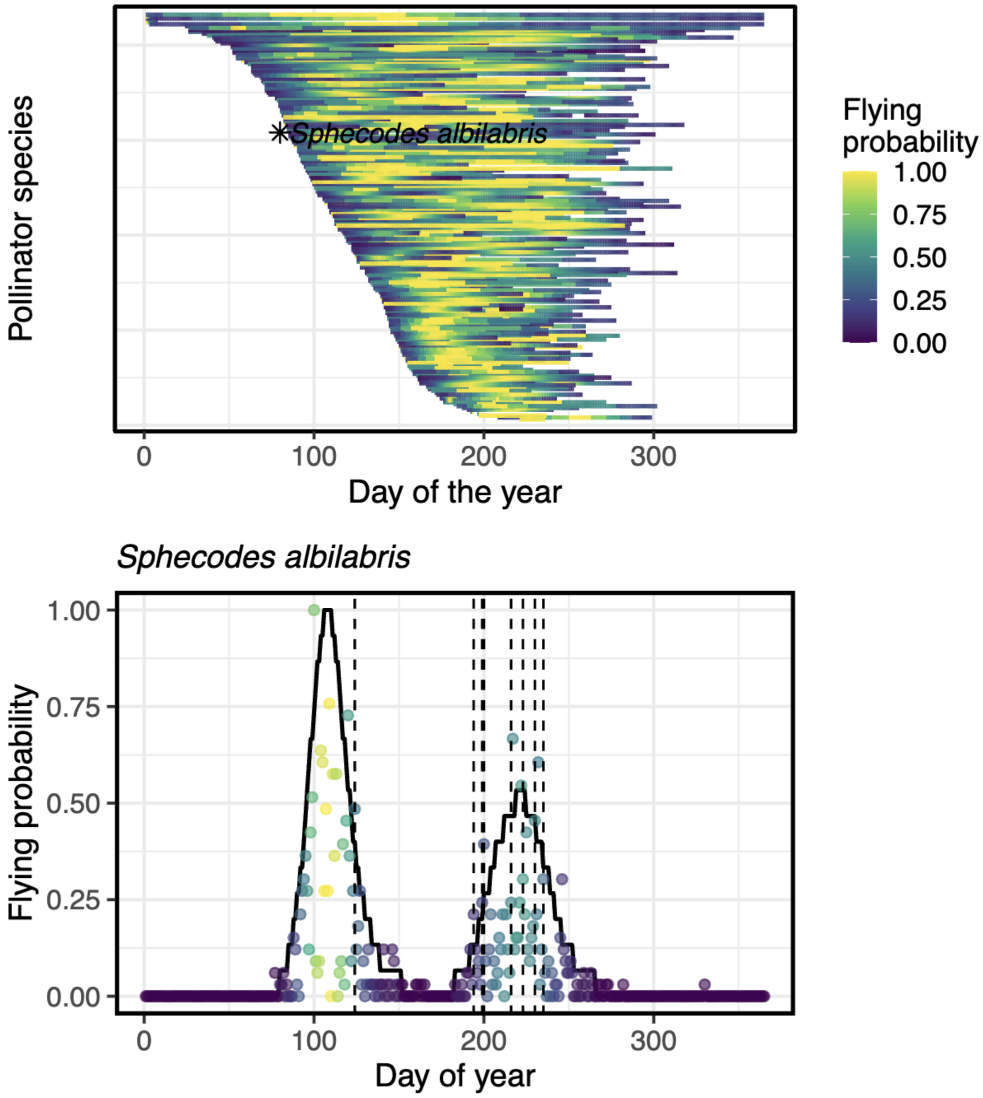


**Figure S5.** Pollinator phenology estimated from GBIF occurrence records. (A) Predicted flying periods for all pollinator species included in the study, ordered by the onset of their flying period. Colour indicates the estimated probability of occurrence (flying probability) across the season. The highlighted species (*Sphecodes albilabris*) is shown as an example. (B) Phenology of the example species, showing the fitted flying probability curve (solid line) together with GBIF occurrence records (points, scaled by relative abundance). Dashed vertical lines indicate days on which the species was observed interacting with focal plants in this study. The close correspondence between observed interactions and predicted flying periods supports the use of GBIF-based occurrence data to estimate pollinator phenology.

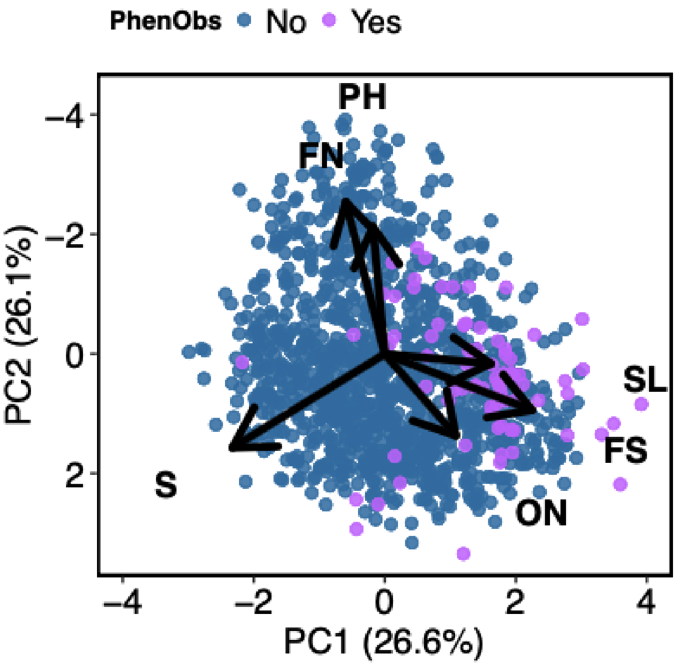


**Figure S6.** Position of focal plant species within the global angiosperm reproductive trait spectrum. Blue points represent species from the global dataset of floral reproductive traits described by Lanuza et al. (2023). Purple points correspond to the PhenObs species monitored in this study. Arrows indicate the loadings of the main reproductive traits on the first two principal components. The focal species cluster within the region of the spectrum associated with higher investment in reproductive traits linked to pollinator attraction, consistent with the focus on pollinator-dependent taxa in this study. This figure illustrates the position of the sampled taxa within the broader reproductive trait space rather than representing a formal analysis.

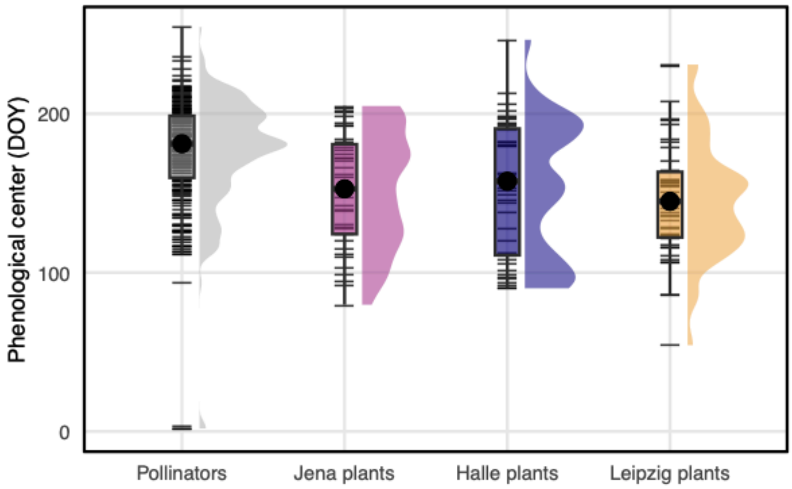


**Figure S7.** Phenological centres of flowering plants and pollinators across botanical gardens. Distributions of phenological centres (day of year, DOY) are shown for pollinators and flowering plants in Jena, Halle and Leipzig. Pollinator phenology was estimated at the regional level from occurrence records and is therefore represented as a single distribution. Half-eye densities and boxplots show the distribution and spread of values, with black points indicating medians and horizontal line segments representing individual observations. Pollinators peaked later in the season than plants.

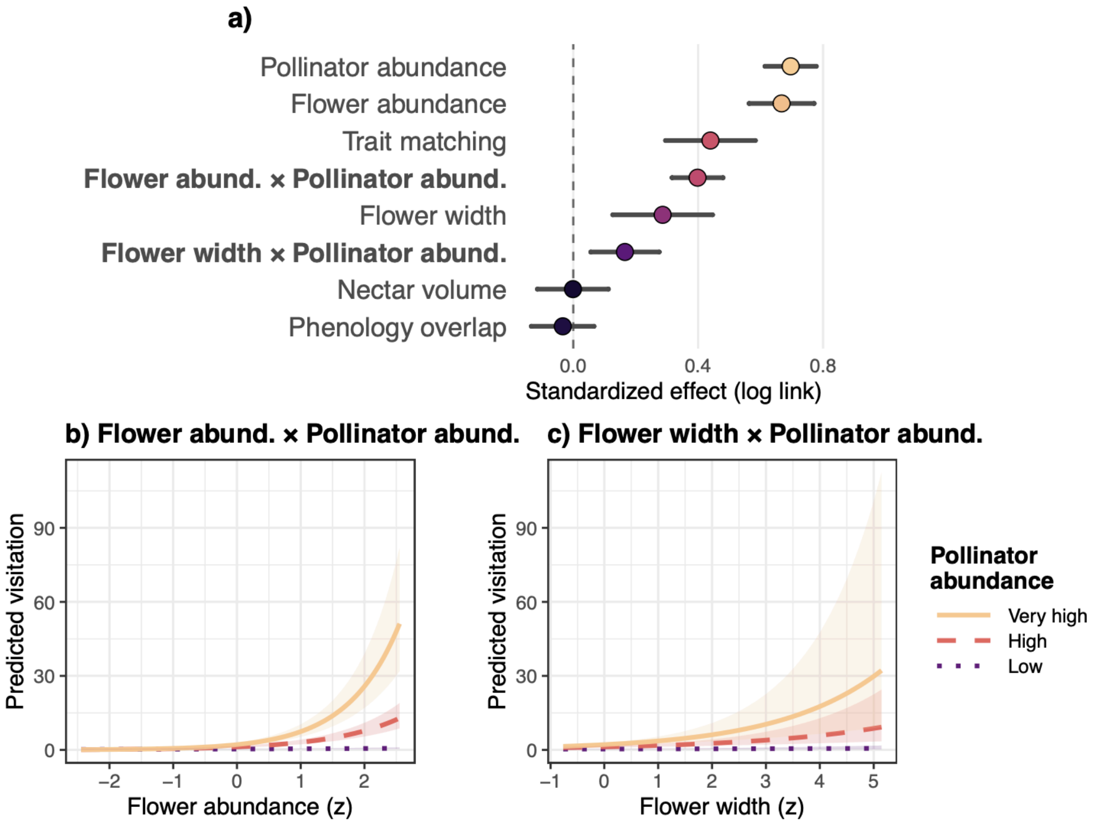


**Figure S8.** Effects of ecological ecological variables (i.e., abundance, traits, and phenology) on the number of plant-pollinator interactions. (a) Standardized model coefficients (log link scale) with 95% confidence intervals, illustrating the direction and relative strength of the ecological variables. (c-b) Predicted number of plant-pollinator interactions across gradients of flower abundance (c) and flower width. Lines show predicted values from the fitted model and shaded areas represent 95% confidence intervals. Different line types and colour gradients indicate pollinator abundance levels corresponding to the 10th (low), 75th (high), and 90th (very high) percentiles of standardized log pollinator abundance.

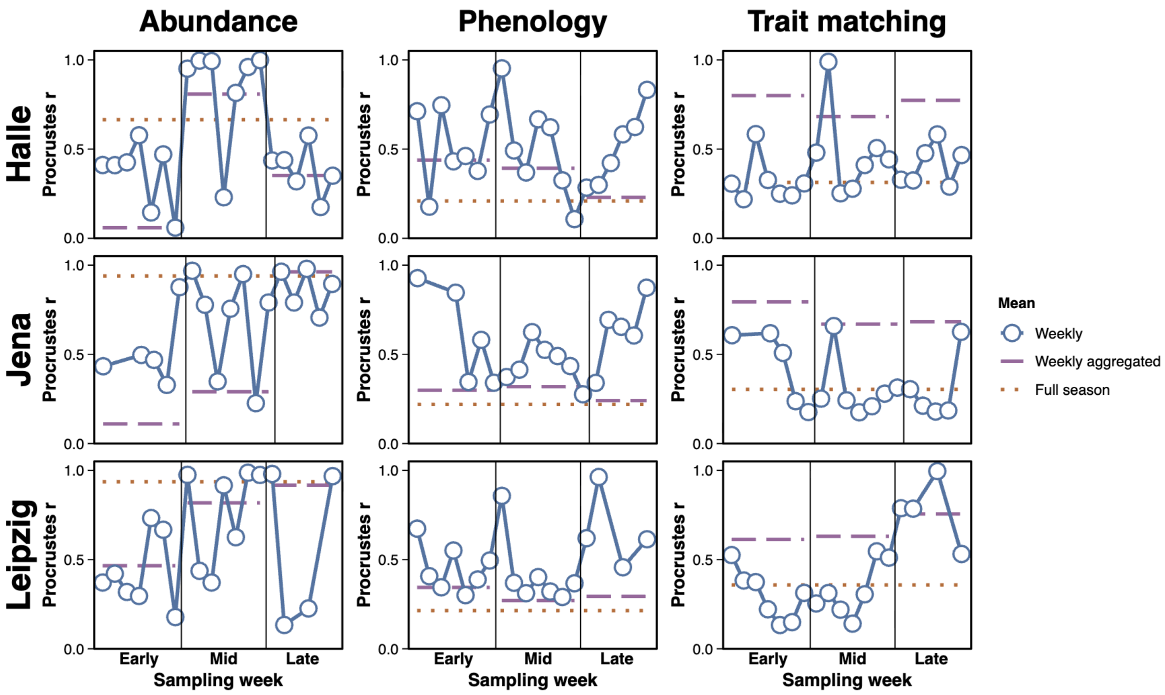


**Figure S9.** Procrustes association between visitation rate and plant-pollinator interaction probability matrices derived from abundance (left column), phenology (middle column), and trait matching (right column) across levels of temporal resolution (weekly, weekly aggregated, and full season). Columns correspond to the three species attributes, whereas rows represent the three botanical gardens (Halle, Jena, and Leipzig). Weekly aggregated values correspond to three seasonal categories (Early, Mid, and Late), obtained by pooling weekly networks into three equal temporal partitions of the full sampling period. Weekly values represent individual weekly networks within each seasonal category. Solid lines and points represent mean Procrustes associations for weekly networks, whereas dashed and dotted horizontal lines indicate weekly aggregated seasonal networks and full-season networks, respectively.

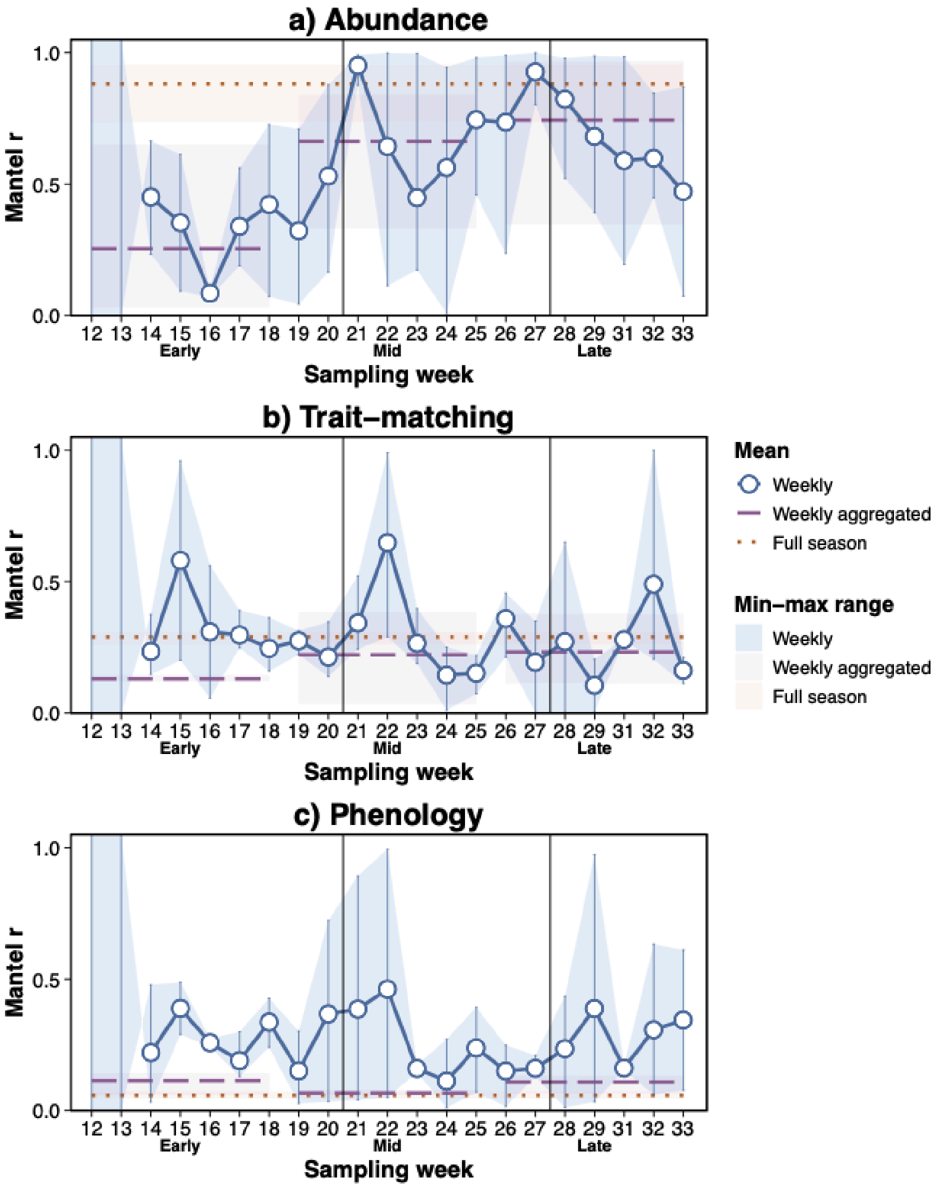


**Figure S10**. Mantel correlation between visitation rate and abundance (a), phenology (b), and traits (size matching; c) across botanical gardens and three levels of temporal complexity (weekly, weekly aggregated and full season). Weekly aggregated values correspond to three seasonal categories (Early, Mid, and Late), obtained by pooling weekly networks into three equal temporal partitions of the full sampling period. Weekly values represent individual weekly networks and are displayed within the corresponding seasonal category. Because multiple weekly values are available per garden and seasonal category, weekly data are shown as individual points together with their mean and 95% confidence intervals (± 1.96 SE). Full-season values correspond to networks aggregated across the entire sampling period for each botanical garden. Temporal complexity is represented by point size, and botanical garden by colour.

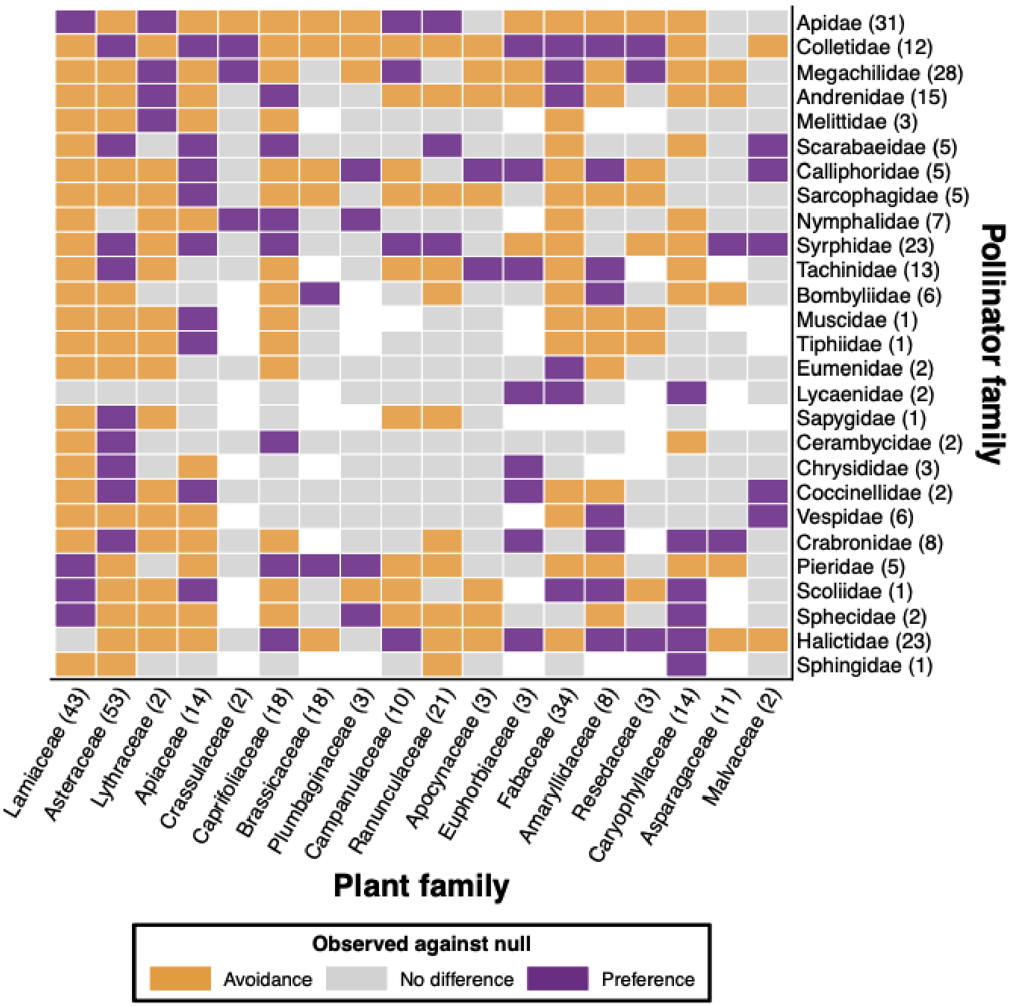


**Figure S11**. Heatmap of standardized effect sizes (SES) of plant-pollinator interactions relative to a spatiotemporally constrained null model at the **plant family × pollinator family** resolution. Colours indicate deviations from null expectations (avoidance, no difference, and preference). Numbers in parentheses indicate species richness for each pollinator family (y axis) and plant family (x axis).

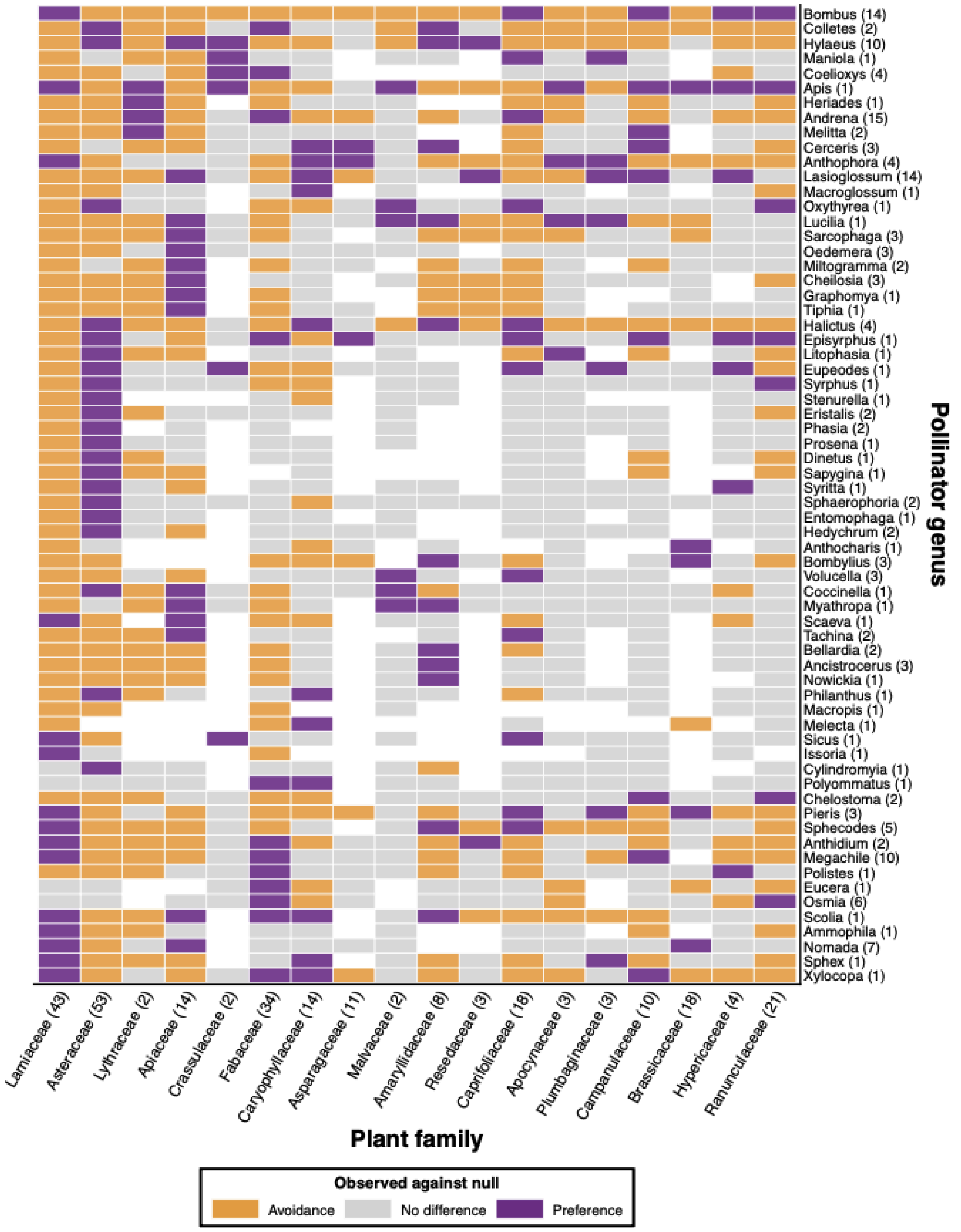


**Figure S12**. Heatmap of standardized effect sizes (SES) of plant-pollinator interactions relative to a spatiotemporally constrained null model at the **plant family × pollinator genus** resolution. Colours indicate deviations from null expectations (avoidance, no difference, and preference). Numbers in parentheses indicate species richness for each pollinator genus (y axis) and plant family (x axis).

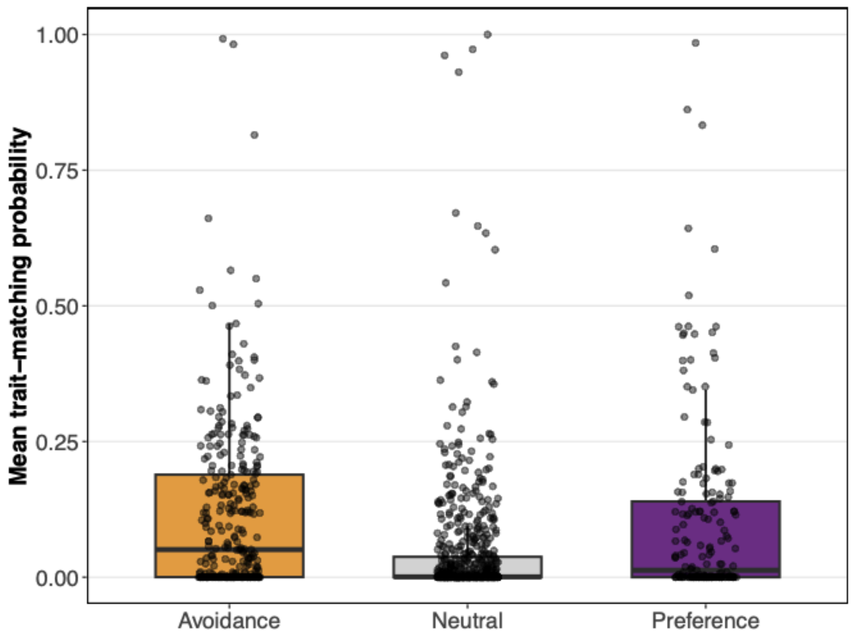


**Figure S13**. Morphological trait matching probability across interaction categories identified by the null-model analysis at the plant family × pollinator genus level. Interaction categories represent significant avoidance (SES < −1.96), no difference from null expectations (−1.96 ≤ SES ≤ 1.96), and significant preference (SES > 1.96).

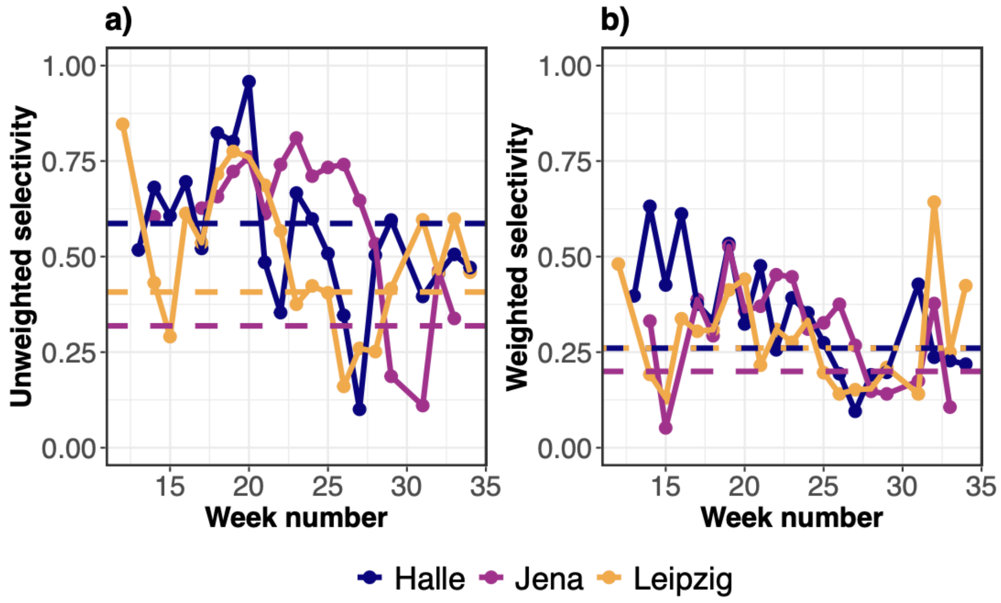


**Figure S14**. Temporal variation in mean pollinator selectivity across botanical gardens. Weekly mean selectivity (d') was calculated from interaction frequency matrices (a) and after accounting for floral resource availability (b).
